# The phosphate exporter XPR1 regulates a gasdermin D–independent mature IL-1β secretion pathway in LPS-stimulated human monocytic cells

**DOI:** 10.64898/2025.12.23.695886

**Authors:** Nathalie Remy, Charlotte Six, Sandrine Tury, Emile Van Schaftingen, Jean-Luc Battini, Sophie Lucas, Pierre G. Coulie, Tiphanie Gomard

## Abstract

The inflammatory cytokine interleukin (IL)-1β is a leaderless protein that is not secreted via the classical endoplasmic reticulum-Golgi pathway but instead is secreted during pyroptosis, a form of caspase-dependent inflammatory cell death mediated by gasdermin D (GSDMD) cleavage and pore formation at the plasma membrane. However, human monocytes can secrete IL-1β in the absence of cell death, and the contribution of GSDMD in this secretory pathway is not established. Here, we identify two mechanisms of mature IL-1β secretion by living human monocytic cells: a rapid, GSDMD-dependent pathway and a slower, GSDMD-independent pathway. Using a CRISPR-Cas9 loss-of-function screen, we identified *XPR1* (Xenotropic and Polytropic retrovirus Receptor 1) as a key regulator of the GSDMD-independent pathway. XPR1 is the only phosphate exporter identified in metazoans and has no previously described function in cytokine secretion. *XPR1* invalidation in *GSDMD*^-/-^ monocytic cells impaired IL-1β secretion. We further show that this regulatory function requires cell-surface expression of XPR1 and is linked to its phosphate export activity. Our results reveal a previously undescribed mechanism of IL-1β secretion by living human monocytic cells, independent of GSDMD and unexpectedly linked to phosphate homeostasis. Deciphering this pathway could lead to new therapeutic modalities for IL-1β-driven inflammatory diseases.

## INTRODUCTION

Interleukin (IL)-1β is an inflammatory cytokine that plays a key role in the early phases of innate immune responses to pathogens^1^. It also is involved in autoinflammatory diseases such as atherosclerosis, rheumatoid arthritis, gout, and type 2 diabetes ^1,2^. IL-1β is secreted mainly by monocytes and macrophages through a tightly regulated multistep process. A first signal provided by microbial or danger-associated molecules induces expression of the *IL1B* gene, which encodes pro-IL-1β (31 kDa), the inactive precursor of IL-1β. The pro-IL-1β N-terminal propeptide then is cleaved by caspase-1 to generate the mature and biologically active IL-1β (17 kDa) in a process that requires prior activation of the inflammasome ^3^. Finally, mature IL-1β, which has no signal peptide, is secreted through unconventional secretory pathways, i.e., independently of the endoplasmic reticulum-Golgi system.

Canonical inflammasomes are composed of sensor proteins such as NLR Family Pyrin Domain-Containing Protein 3 and NLR Family CARD Domain-Containing Protein 4. Microbial or sterile cellular stress signals such as lipopolysaccharide (LPS), the bacterial toxin nigericin, and extracellular ATP activate NLRP3 ^4–6^, which recruits and forms pro-caspase-1 filaments, leading to activated caspase-1 ^3,7^. The latter cleaves pro-IL-1β and also gasdermin D (GSDMD), releasing the C-terminal repressor domain of GSDMD from its p30 N-terminal pore-forming domain ^8^. The pore-forming domain binds with high affinity to negatively charged lipids restricted to the inner leaflet of the plasma membrane ^9^, then oligomerizes to form pores with an inner diameter of about 20 nm ^10^. The resulting cell swelling can be followed by plasma membrane rupture and cell death, accompanied by the release of intracellular proinflammatory mediators, including mature IL-1β. This inflammatory cell death program is called pyroptosis.

In 2015, GSDMD-dependent pyroptotic cell death was shown to be associated with a release of mature IL-1β ^11–13^, leading to the widespread concept that IL-1β secretion requires cell death. However, several groups have observed IL-1β secretion in the absence of cell death, including in murine neutrophils during acute *Salmonella* challenge ^14,15^, murine macrophages activated by peptidoglycan ^16^, murine dendritic cells primed with LPS and treated with oxidized 1-palmitoyl-2-arachidonoyl-sn-glycero-3-phosphorylcholine ^17^, and human monocytes stimulated with LPS ^18,19^. Like other groups, we observed that primary human monocytes stimulated with LPS secreted mature IL-1β without detectable cell death. How these living cells secrete mature IL-1β is not completely understood, although several pathways have been proposed ^20^. A major factor to be considered is the involvement of GSDMD. Indeed, in a model of hyperactivated murine macrophages, mature IL-1β was shown to be secreted through a GSDMD-dependent pathway in the absence of pyroptosis ^21^.

Here we evaluated the putative GSDMD requirement for mature IL-1β secretion by living human monocytic cells. Using a cellular model that allows for gene editing, we demonstrated the existence of an IL-1β secretion pathway that is independent of GSDMD and requires the phosphate exporter Xenotropic and Polytropic retrovirus Receptor 1 (XPR1).

## RESULTS

### THP-1 cells stimulated with LPS secrete mature IL-1β without undergoing pyroptosis

To study the mechanism of mature IL-1β secretion by human monocytic cells, we used the spontaneously immortalized cell line THP-1, derived from a patient with acute monocytic leukemia ^22^. The mechanisms underlying the secretion of mature IL-1β by THP-1 cells stimulated with LPS alone have not been thoroughly investigated, and several authors have suggested that THP-1 cells may not constitute an appropriate model for such studies ^18^. The response of THP-1 to LPS is typically studied following treatment with PMA, which induces their differentiation into adherent, macrophage-like cells^23^. Among the various differentiation protocols reported in the literature, we selected a treatment with 300 ng/mL PMA for 3h, followed by an 18 h rest period in PMA-free medium prior to stimulation with 10 ng/mL LPS, without the addition of any inflammasome activators^24^ (Fig. 1A, Fig. S1). Under these conditions, mature IL-1β was detected in the supernatants after 24 h, as measured by ELISA (Fig. 1B). This secretion was not accompanied by cell death, as indicated by the absence of a significant increase in lactate dehydrogenase (LDH) release, the low proportion of cells stained with the small DNA intercalant CellTox Green, and the lack of a significant decrease in cellular metabolic activity assessed with a tetrazolium reduction assay (Fig. 1B). Thus, upon LPS stimulation, PMA-differentiated THP-1 cells secrete mature IL-1β without undergoing pyroptosis or another form of cell death. Using knockout cells, we showed that this LPS-induced secretion of mature IL-1β is dependent on NLRP3 and Caspase-1, but not on caspase-4 or caspase-5 (non-canonical inflammasome pathway^25^), nor on caspase-8 (alternative inflammasome pathway^18^) (Fig S2). By contrast, addition of the bacterial toxin nigericin 4 h after LPS stimulation markedly enhanced mIL-1β secretion-by approximatively sevenfold- and triggered robust pyroptotic cell death, as evidenced by an increased proportion of CellTox Green–positive cells, elevated LDH release and a 20-fold decrease in metabolic activity (Fig. 1B). Together, these results indicate that PMA-differentiated THP-1 cells recapitulate the distinctive ability of primary human monocytes to secrete mature IL-1β in response to LPS alone, without undergoing pyroptosis.

**Figure 1.**
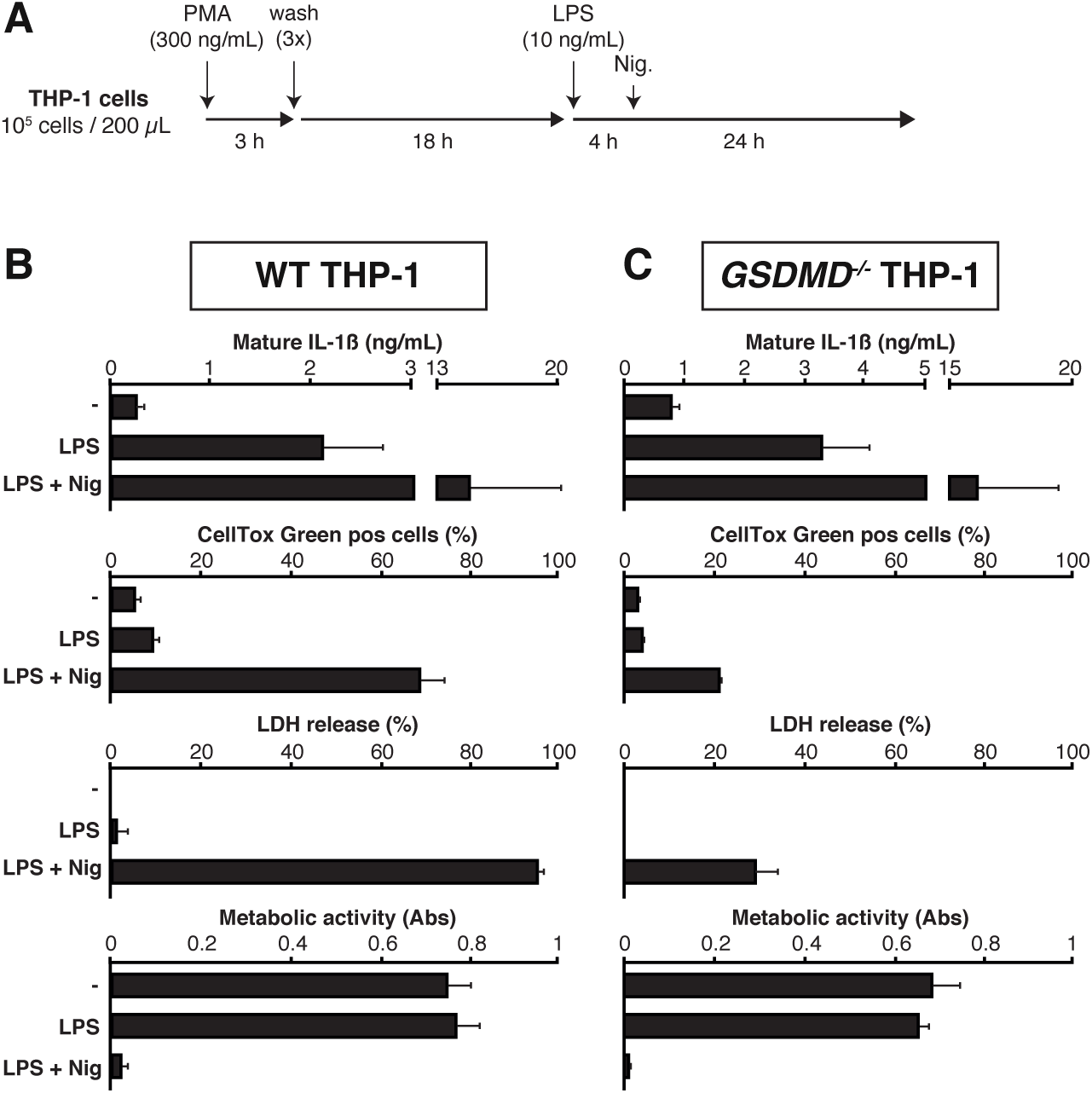
PMA-differentiated THP-1 cells secrete mature IL-1β in the absence of cell death and independently of GSDMD. (A) Experimental workflow of PMA differentiation and LPS stimulation of THP-1 cells. Cells were seeded at 1 x 10^5^ cells per well (200 µL) and differentiated with PMA (300ng/mL) for 3 h. After three washes, cells were incubated in PMA-free medium for 18 h. Cells were then stimulated with 10 ng/mL ultrapure LPS from *E. coli* for 24 h. As a positive control for cell death, pyroptosis was induced by nigericin (Nig.) (10 µM) added 4 h after LPS. See also Figure S1. (B) Secretion of mature IL-1β and assessment of cell viability following LPS stimulation of wild-type (WT) or (C) *GSDMD*^-/-^ THP-1 cells. PMA-differentiated THP-1 cells (WT of *GSDMD*^-/-^) were stimulated with LPS (10 ng/mL) or not for 24 h. Secreted mature IL-1β was measured by ELISA. Cell death was evaluated by CellTox^TM^ Green uptake and LDH release. Metabolic activity was assessed using a MTS/PMS assay. Nigericin (10µM), added 4 h after LPS, was used as a positive control for pyroptosis. Data are shown as means ± SEM of biological triplicates. Each panel is representative of at least three independent experiments. The results with *GSDMD*^-/-^THP-1 cells were reproduced with two different clones generated with distinct gRNAs (Table S1), and complete gene invalidation was verified by targeted sequencing (Table S1).

### Mature IL-1β secretion by PMA-treated, LPS-stimulated THP-1 cells does not require GSDMD

To assess the involvement of GSDMD in the secretion of mature IL-1β by THP-1 cells, we engineered GSDMD^-/-^ THP-1 clones with CRISPR/Cas9 and validated them by sequencing (Table S1). Surprisingly, GSDMD^-/-^ THP-1 clones stimulated with LPS did not secrete less mature IL-1β than wild-type (WT) THP-1 cells, revealing the existence of a GSDMD-independent secretory pathway (Fig. 1C). Similar to WT THP1 cells, the GSDMD^-/-^ THP-1 clones stimulated with LPS secreted mature IL-1β without undergoing cell death (Fig. 1C).

Although the amounts of secreted mature IL-1β were the same for WT and GSDMD^-/-^ THP-1 cells after 24 h of LPS stimulation, the kinetics of the secretion were strikingly different (Fig. 2A). In WT cells, mature IL-1β secretion was rapid, with a sharp increase between 2 and 4 h after LPS addition, and contrasted with that of the cytokine Interleukin-1 Receptor Antagonist (IL-1RA), secreted by the conventional endoplasmic reticulum–Golgi pathway (Fig. 2B). In contrast, GSDMD^-/-^ THP-1 cells secreted mature IL-1β at a slower rate than WT cells, with minimal secretion during the first 8 h following LPS stimulation (Fig. 2A).

**Figure 2.**
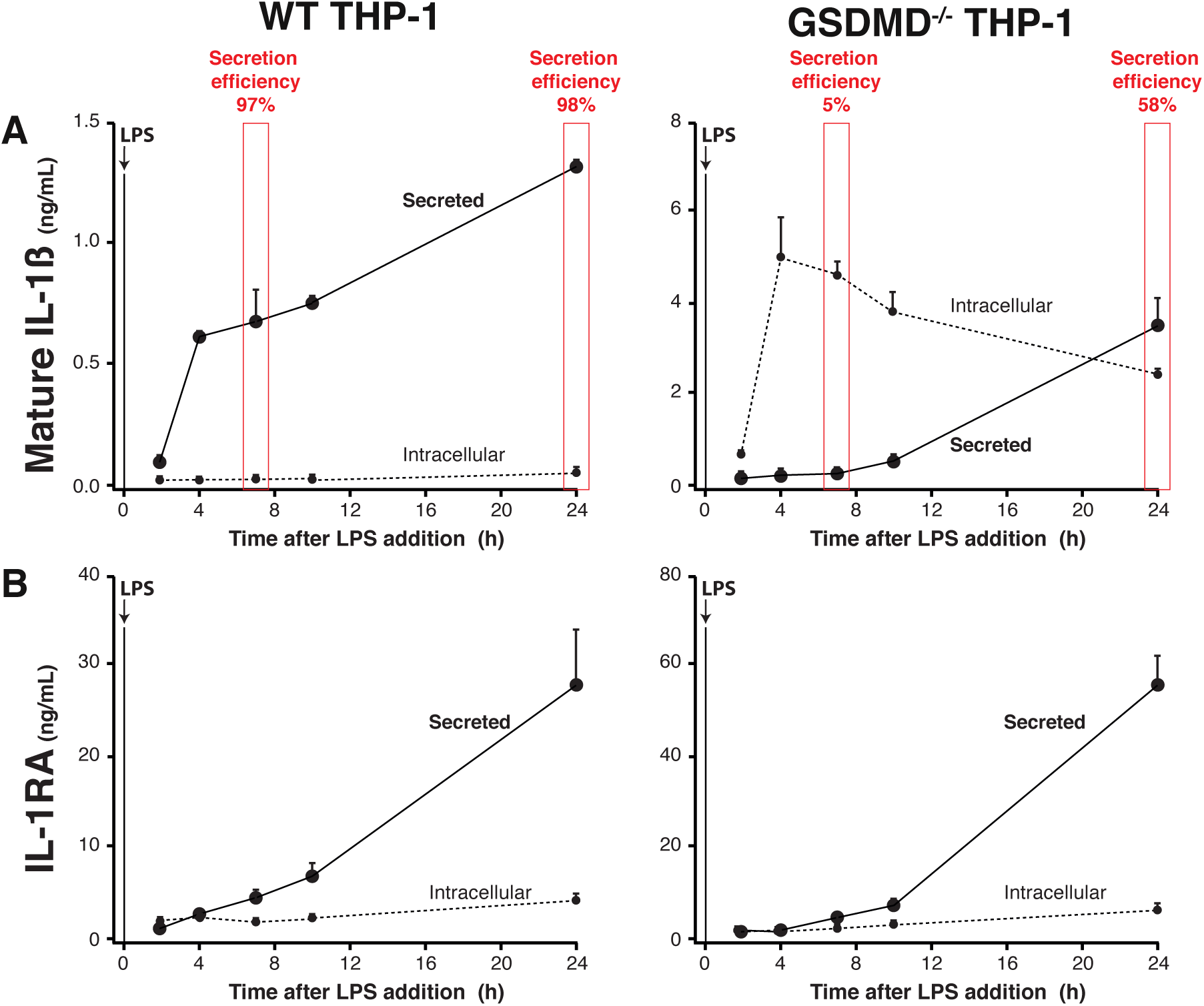
PMA-differentiated and LPS-stimulated THP-1 cells secrete mature IL-1β through GSDMD-dependent and -independent pathways. WT or *GSDMD*^-/-^ PMA-differentiated THP-1 cells were stimulated with LPS (10 ng/mL) for 24 h. At 2, 4, 7, 10, and 24 h, supernatants were harvested and cells were resuspended in 200 µL of fresh medium (same volume as supernatants). Cells and supernatants were frozen at -20 °C for at least 18 h. After thawing, cell lysates were clarified by centrifugation to remove cellular debris. Mature IL-1β (A) and IL-1RA (B) were measured by ELISA in both supernatants and cell lysates. Secretion efficiency after 7 or 24 h of LPS stimulation was calculated as the ratio of the cytokine content measured in supernatant (200µL) to the total cytokine content measured in the supernatant plus the corresponding cell lysate (200µL). The results with *GSDMD*^-/-^ THP-1 cells were reproduced with two different clones generated with distinct gRNAs (Table S1), and complete gene invalidation was verified by targeted sequencing (Table S1). See also Figure S3.

GSDMD is not known to play a role in the production of mature IL-1β, which depends on *IL1B* gene transcription and cleavage of pro-IL-1β into mature IL-1β. Therefore, both the amounts and the rate of production of mature IL-1β were expected to be similar in WT and GSDMD^-/-^ THP-1 cells. We reasoned that the slower rate of secretion by GSDMD^-/-^ THP-1 cells could be associated with intracellular accumulation of the cytokine. To test this, we measured the concentrations of mature IL-1β in cell pellets lysed by a freeze–thaw cycle. In WT THP-1 cells, no intracellular mature IL-1β could be detected (Fig. 2A). In contrast, in GSDMD^-/-^ THP-1 cells, large amounts of intracellular mature IL-1β were measured 4 h after LPS addition, slowly decreasing as secretion occurred. These results, obtained by ELISA on cell pellets, were confirmed by immunoblot analysis (Fig S3).

We calculated the efficiency of mature IL-1β secretion as the proportion of total (intracellular + secreted) mature IL-1β found in cell supernatants. In WT THP-1 cells, secretion efficiency was consistently high, reaching 97% and 98 % at 7 and 24 hours after LPS addition, respectively (Fig. 2A). In GSDMD^-/-^ THP-1 cells, secretion efficiency was very low (5%) at 7 hours after LPS addition and increased up to 58% by 24 hours after LPS stimulation (Fig. 2A). Notably, no intracellular accumulation of IL-1RA was observed in the GSDMD^-/-^ THP-1 cells (Fig. 2B), indicating that the absence of GSDMD did not impair the efficiency of the conventional ER-Golgi-dependent secretion pathway.

We conclude that THP-1 cells stimulated with LPS alone secrete mature IL-1β in a cell death–independent manner via at least two distinct pathways. The first pathway is GSDMD-dependent and mediates rapid and efficient cytokine secretion without intracellular accumulation. The second pathway is GSDMD-independent and results in slower and less efficient secretion.

### XPR1 is involved in the GSDMD-independent pathway of mature IL-1β secretion by LPS-stimulated THP-1 cells

To identify proteins involved in GSDMD-independent IL-1β secretion, we used the genome-wide CRISPR Brunello library in GSDMD^-/-^ THP-1 cells. Transduced cells were differentiated with PMA, stimulated with LPS for 24 h, labeled for intracellular mature IL-1β, and sorted by flow cytometry to isolate the cells with the highest content of mature IL-1β (Fig. 3A and Fig. S4). We hypothesized that the cells that accumulated the most mature IL-1β would correspond to cells in which a guide (g)RNA invalidated a gene involved in the GSDMD-independent secretion pathway of mature IL-1β. One of the most enriched gene hits in these cells was *XPR1*, first described as a retroviral receptor ^26–28^ (Fig. 3B, Dataset S1 and S2). XPR1 is a ubiquitously expressed multipass surface membrane protein and, to date, the only known phosphate exporter in metazoans ^29^. We verified that XPR1 was expressed on WT and GSDMD^-/-^ THP-1 cells (Fig. 3C).

**Figure 3.**
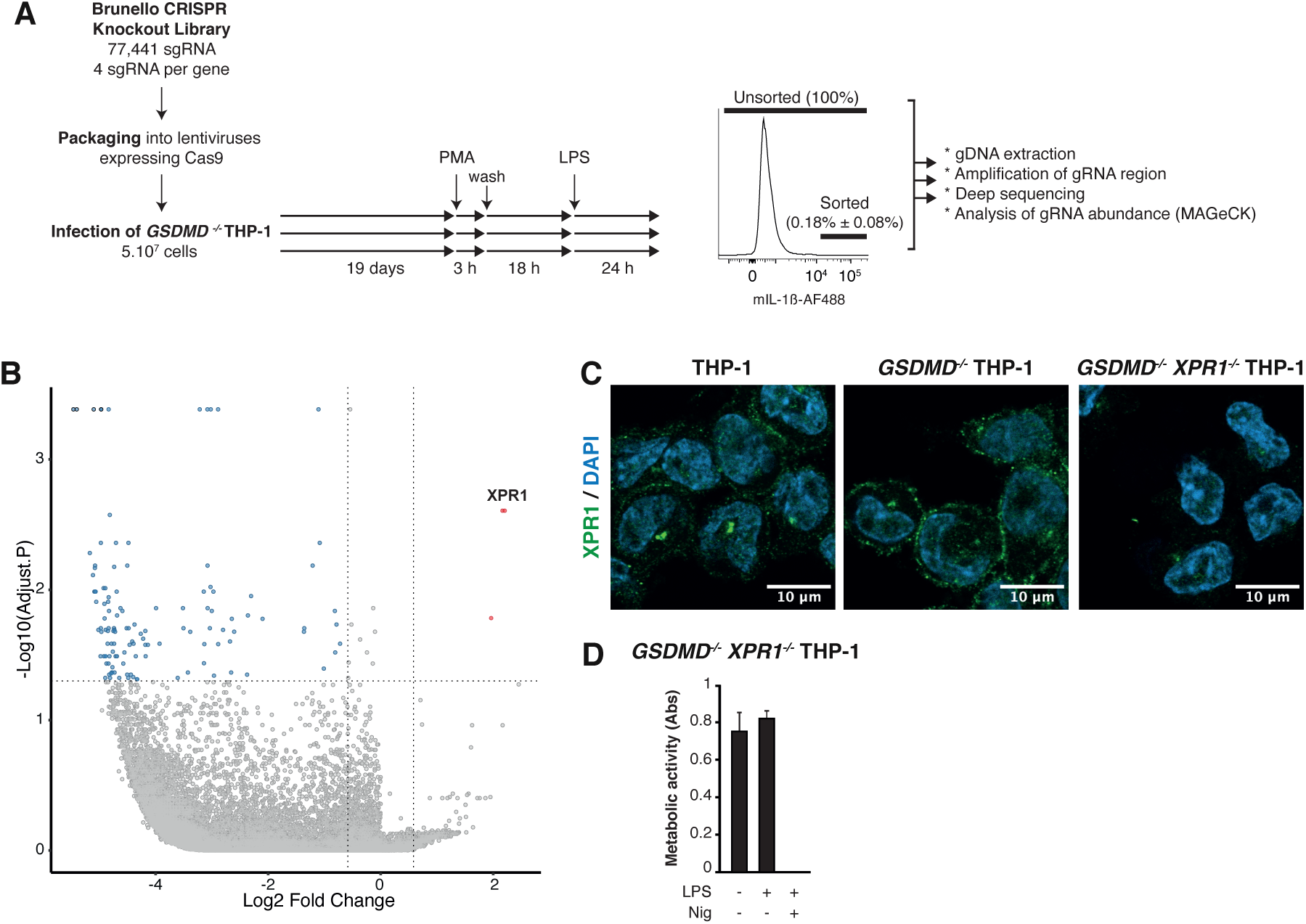
Genome-wide CRISPR screening identifies XPR1 as a regulator of GSDMD-independent IL-1β secretion. (A) Schematic overview of the CRISPR/Cas9 library screening strategy. The genome-wide human library Brunello, encoding Cas9 and 77,441 single guide (sg)RNAs, was packaged into lentiviruses produced by HEK293 cells and immediately used to transduce *GSDMD*^-/-^ THP-1 cells. Three independent batches of THP-1 cells were infected on the same day with the same pool of viruses. After 19 days of culture, transduced cells were differentiated with PMA and stimulated with LPS for 24 h. After intracellular labeling of mature IL-1β, the cells of each batch displaying the highest staining were sorted by flow cytometry. Genomic DNA of sorted cells, together with control DNA from unselected transduced cells, was used as a template for sgRNA PCR amplification, followed by high-throughput sequencing. sgRNA abundance and enrichment were analyzed with MAGeCK. See also Dataset S1 and S2 and Figure S4. (B) Volcano plot showing gene enrichment and depletion in the screen. For each gene, the x axis shows its enrichment or depletion, calculated as the mean of all four sgRNAs targeting the gene, in the three sorted mIL-1β positive populations relative to the corresponding unsorted populations (paired analysis). The y axis shows statistical significance. Significantly enriched genes are highlighted in red. (C) Representative immunofluorescence images of WT, *GSDMD*^-/-^, and *GSDMD*^-/-^ *XPR1*^-/-^ THP-1 cells differentiated with PMA and labeled with anti-XPR1 (green) and DAPI (blue). Magnification 63×; scale bar, 10 µm. (D) Cell viability following LPS stimulation of XPR1-invalidated THP-1 cells. *GSDMD*^-/-^ *XPR1*^-/-^THP-1 cells were differentiated with PMA and stimulated or not with LPS for 24 h. Metabolic activity was assessed with the MTS/PMS assay. As a positive control for cell death, pyroptosis was induced by addition of nigericin (Nig) (10 µM) 4 h after LPS stimulation. Data are represented as means ± SEM of biological triplicates. Data shown are representative of at least three independent experiments. The results shown for gene-invalidated cells in panels C and D were reproduced with two different clones generated with distinct gRNAs (Table S1), and complete gene invalidation was verified by targeted sequencing (Table S1).

We then generated *GSDMD*^-/-^ *XPR1*^-/-^ double-knockout THP-1 cells, which remained viable and metabolically active after differentiation with PMA and stimulation with LPS (Fig. 3D and Table S1). PMA-differentiated and LPS-stimulated *GSDMD*^-/-^ *XPR1*^-/-^ cells displayed a significantly lower secretion efficiency of mature IL-1β than the *GSDMD*^-/-^ THP-1 cells (19% vs 58%; *P* < 0.0001), confirming the involvement of XPR1 in the GSDMD-independent secretion of mature IL-1β under these conditions (Fig. 4A and Fig. S5A).

**Figure 4.**
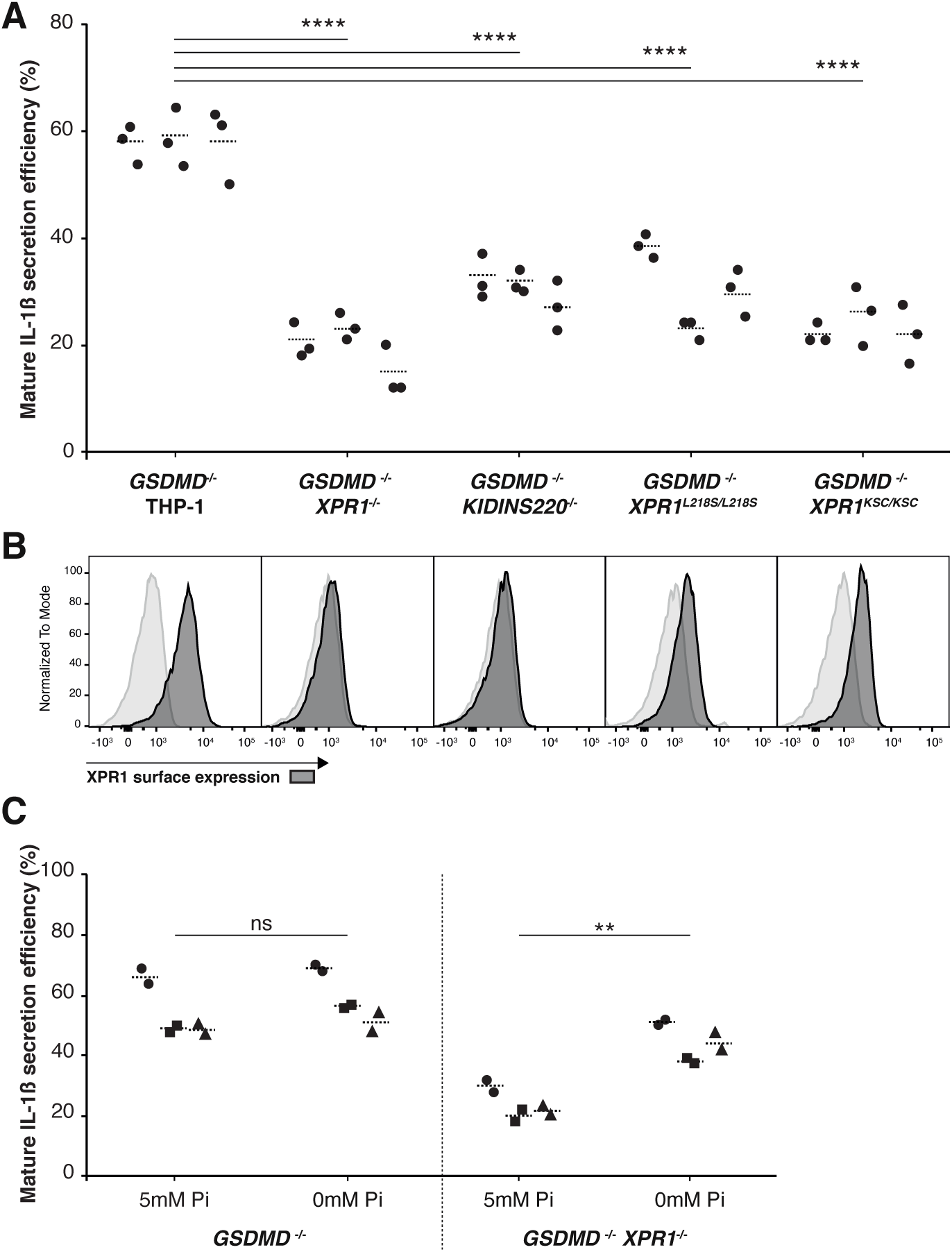
XPR1 positively regulates GSDMD-independent secretion efficiency of mature IL-1β potentially through its phosphate export function. (A) Mature IL-1β secretion efficiencies by indicated gene-edited THP-1 cells after 24 h of LPS stimulation, measured and calculated as indicated in the legend of Fig. 2. Data represent three biological replicates from each of three independent experiments. Statistical analysis was performed using a nested one-way analysis of variance with Tukey’s test; ****, *P* ≤ 0.0001; all other comparisons were not significant. (B) Cell-surface expression of XPR1. Cells were labeled with an anti-XPR1 immunoadhesin ligand comprising the receptor–binding domain of the xenotropic murine leukemia virus envelope glycoprotein (XRBD) fused to mouse IgG1 Fc, followed by a PE-conjugated anti-mouse IgG1 secondary antibody. Results were reproduced with two different clones generated with distinct gRNAs (Tables S1 and S2). For each clone, complete gene invalidation was verified by targeted sequencing (Tables S1 and S2). See also Figure S5. (C) Mature IL-1β secretion efficiencies by the indicated gene-edited THP-1 cells after PMA-differentiation and 24 h of LPS stimulation in RPMI without phosphate (0 mM Pi) or supplemented with 5mM Pi, measured and calculated as described in the legend of Fig. 2. Data represent two biological replicates from each of three independent experiments. Statistical analysis was performed using a nested one-way analysis of variance with Tukey’s test; **, *P* ≤ 0.01; ns, nonsignificant; Pi, inorganic phosphate.

It is now well established that XPR1 forms a complex with KIDINS220 (Kinase D-interacting substrate of 220 kDa), a cell surface protein with four transmembrane domains, and that this interaction is required for proper surface localization and phosphate-export activity of XPR1^30,31^. To further establish evidence of the involvement of XPR1 in mature IL-1β secretion, we produced *GSDMD*^-/-^ *KIDINS220*^-/-^ double-knockout THP-1 cells (Table S1). We confirmed that these cells had no detectable surface expression of XPR1 (Fig. 4B and Fig. S5B). Like the *GSDMD*^-/-^ *XPR1*^-/-^THP-1 cells, the *GSDMD*^-/-^ *KIDINS220*^-/-^ THP-1 double knockouts secreted mature IL-1β less efficiently than their *GSDMD*^-/-^ THP-1 counterparts (30% vs 58%; *P* < 0.0001), suggesting that cell surface expression of XPR1 is required for its role in mature IL-1β secretion (Fig. 4A and Fig. S5A). Altogether, these results indicate that XPR1 and KIDINS220 are necessary for GSDMD-independent secretion of mature IL-1β by THP-1 cells.

### XPR1 positively regulates GSDMD-independent IL-1β secretion through regulation of intracellular phosphate levels

XPR1 is a 696 residue multi-transmembrane protein with both intracellular amino and carboxy termini. The aminoterminal region contains a 177-residue domain called SPX (for SYG1, Pho81 and XPR1: yeast and human SPX-containing proteins), found in many yeast and plant proteins directly involved in phosphate homeostasis or regulated themselves by phosphate levels. The SPX domain is crucial for XPR1 phosphate export function. Indeed, many XPR1 pathogenic variants, identified in patients with a neurodegenerative disease called primary familial brain calcifications (PFBC) ^32^, are mostly located within or next to this SPX domain and alter phosphate export (for instance the L218S PBFC mutation). Moreover, the SPX domain harbors a “lysine surface cluster” (KSC), that delineates a binding pocket for inositol pyrophosphate, more specifically diphospho-*myo*-inositol pentakisphosphate (5-InsP_7_) and bis-diphospho-*myo*-inositol tetrakisphosphate (1,5-InsP_8_), which are signaling molecules in high phosphate conditions ^33^. Mutations in this KSC pocket (triple mutant K158A/K161A/K165A) altered phosphate efflux, revealing the importance of inositol pyrophosphate binding to the SPX domain of XPR1 for efficient phosphate efflux function ^34–36^.

To investigate the role of XPR1-mediated phosphate export in GSDMD-independent mature IL-1β secretion, we engineered *GSDMD*^-/-^ *XPR1^L218S/L218S^*and *GSDMD*^-/-^ *XPR1^KSC/KSC^* THP-1 cells by genome editing of the two endogenous *XPR1* copies of THP-1 cells using CRISPR-mediated homology-directed repair (Table S2). This strategy circumvents the artifact associated with overexpression of transduced mutant *XPR1* transgenes. Cell surface expression of WT and mutant XPR1 proteins was verified by labeling with a well-characterized XPR1 ligand ^29^ (Fig. 4B and Fig. S3B). Using this sensitive assay, WT XPR1 was not detected on the surface of the *GSDMD*^-/-^ *XPR1*^-/-^ or *GSDMD*^-/-^ *KIDINS220*^-/-^ cells. In contrast, mutant XPR1 proteins were easily detected on the *GSDMD*^-/-^ *XPR1^L218S/L218S^* and *GSDMD*^-/-^ *XPR1^KSC/KSC^*cells, although at lower levels than on *GSDMD*^-/-^ cells expressing WT *XPR1* alleles (Fig. 4B). After 24 h of LPS stimulation, the secretion efficiencies of mature IL-1β by *GSDMD*^-/-^ *XPR1^L218S/L218S^* and *GSDMD*^-/-^*XPR1^KSC/KSC^*THP-1 cells were comparable to that of *GSDMD*^-/-^ *XPR1*^-/-^ THP-1 cells, i.e., significantly reduced in comparison with *GSDMD*^-/-^ THP-1 cells (Fig. 4A and Fig. S5A). Although we cannot formally exclude that the reduced surface expression of mutant XPR1 contributes to the lower IL-1β secretion, our results support a role for XPR1-mediated phosphate export in the GSDMD-independent secretion of mature IL-1β by THP-1 cells.

To document a link between phosphate homeostasis and IL-1β secretion, we modified the concentration of inorganic phosphate in the culture medium. When *GSDMD*^-/-^ *XPR1*^-/-^ THP-1 cells were deprived of inorganic phosphate (0 mM Pi versus 5 mM Pi in control medium) during PMA-induced differentiation and LPS stimulation, secretion of mature IL-1β increased significantly (Fig 4C). This effect was specific to XPR1-deficient cells, as it was not observed in *GSDMD*^-/-^ THP-1 cells. These results confirm that inorganic phosphate contributes to the regulation of mature IL-1β secretion.

## DISCUSSION

Our results reveal two previously unrecognized features of mature IL-1β secretion by living monocytic cells. First, we identify two distinct secretory pathways: a rapid, GSDMD-dependent pathway and a slower GSDMD-independent pathway. Second, we demonstrate that the GSDMD-independent pathway requires XPR1, the only known phosphate exporter in metazoans. These findings provide a basis for further investigation into the role of XPR1 in cytokine secretion.

The THP-1 cell line proved to be a suitable model for studying unconventional IL-1β secretion in the absence of pyroptosis. Using GSDMD-deficient THP-1 cells, we provide clear evidence that mature IL-1β can be secreted independently of GSDMD by live human monocytic cells (Fig. 1B-C), consistent with previous observations by Rubartelli et al. of IL-1β release without cell death, propidium iodide uptake or GSDMD cleavage ^19^.

We observed these two IL-1β secretion pathways by tracking the accumulation of mature IL-1β in both culture supernatants and cells over 24h, in the presence or absence of GSDMD. In GSDMD-expressing cells, newly produced IL-1β was rapidly secreted, whereas in GSDMD-deficient cells it accumulated intracellularly before release (Fig. 2A). These results indicate that conclusions about the role of GSDMD in IL-1β secretion may be strongly influenced by the timing of IL-1β measurements.

Considering that we did not observe pyroptosis, we were surprised to detect a GSDMD-dependent component in IL-1β secretion. Mature IL-1β can exit cells through GSDMD pores at the plasma membrane ^10,37^, and GSDMD pores at the plasma membrane can be closed or repaired ^38^. A transient secretion of mature IL-1β through GSDMD pores at the plasma membrane of living cells is therefore possible. However, we consistently failed to detect entry of the small DNA intercalant CellTox Green during the GSDMD-dependent secretion of IL-1β (Fig. 1C). This observation suggests an absence of GSDMD pores at the cell surface and points instead to intracellular GSDMD activity. We are probably far from understanding all of the roles of GSDMD in the cell, but it is known to form pores in the outer and inner membranes of mitochondria ^39^. More work is warranted to elucidate the various functions of GSDMD in mature IL-1β secretion by THP-1 cells.

Here we focused on the GSDMD-independent secretory pathway of mature IL-1β and revealed the involvement of the phosphate exporter XPR1 in this pathway. We obtained this result through an unbiased approach with a genomic CRISPR/Cas9 library (Fig. 3), taking advantage of the transient accumulation of mature IL-1β in the GSDMD^-/-^ THP-1 cells, which peaked at 4 h after LPS addition and then declined. We reasoned that decreasing the secretion should extend the accumulation and lead to IL-1β detectability at 24 h after LPS addition, using fluorescent labeling of intracellular mature IL-1β. In the DNA sequencing data for labeled cells, the MAGeCK algorithm identified *XPR1* as a top candidate in three out of three replicate experiments. We confirmed this surprising result with LPS stimulation of *GSDMD^-/-^ XPR1^-/-^*cells, with a decreased efficiency of secretion of mature IL-1β (Fig. 4).

Recent structural biology studies, mainly using cryo-EM, have revealed major insights into the architecture and regulation of XPR1 ^31,40–46^. These structural analyses show that XPR1 forms a dimeric membrane transporter composed of ten transmembrane helices per monomer organized into an outer scaffold domain and an inner transport core that forms the phosphate-conducting tunnel. The maximal diameter of this tunnel reaches approximately 6-8 Å, a diameter sufficient for inorganic phosphate ions but incompatible with the passage of macromolecules such as mature IL-1β. Thus, any link between XPR1 and IL-1β secretion is unlikely to involve direct transport of the cytokine itself. Rather, given XPR1 is the sole known phosphate exporter in human cells ^29^, alterations in phosphate homeostasis represents a more plausible mechanistic connection.

Phosphate homeostasis is highly complex, and it is remarkable that our CRISPR/Cas9 screening singled out *XPR1*. One possible explanation is the absence of functional redundancy in phosphate export, making XPR1 a critical bottleneck whose disruption directly perturbs intracellular phosphate levels.

Intracellular phosphate homeostasis is maintained by phosphate importers, which in human monocytes are SLC20A1/PiT1 and SLC20A2/PiT2, and by the phosphate exporter XPR1^47^. The impaired secretion of mature IL-1β observed in *GSDMD* ^-/-^ *XPR1* ^-/-^ THP-1 cells is closely linked to defective phosphate homeostasis in these cells. First, we showed that *GSDMD^-/-^* THP-1 cells expressing XPR1 mutants known to be deficient in phosphate efflux display impaired mature IL-1β secretion, mirroring XPR1-deficient cells (Fig. 4A). Second, under phosphate-deprivation conditions for 24 hours, mature IL-1β secretion was increased in the *GSDMD* ^-/-^ *XPR1* ^-/-^ THP-1 cells (Fig. 4C). Although secretion was not fully restored to levels observed in *GSDMD* ^-/-^ THP-1 cells – likely due to residual inorganic phosphate present in serum-supplemented medium- these findings collectively reveal an unexpected role for phosphate homeostasis in the regulation of mature IL-1β secretion.

Inositol pyrophosphates represent a plausible mechanistic link between intracellular phosphate availability and IL-1β secretion (Fig. 5). These highly phosphorylated signaling molecules couple cellular phosphate status to downstream biological processes, and their synthesis is regulated by intracellular inorganic phosphate levels. Indeed, elevated inorganic phosphate stimulates the activity of inositol hexakiphosphate kinases (IP6Ks) diphosphoinositol pentakiphosphate kinases (PPIP5Ks), thereby promoting 5-Insp7 and 1,5-InsP_8_ synthesis ^48^. On the contrary, Pi limitation reduces inositol pyrophosphate synthesis. In turn, inositol pyrophosphate function as sensitive indicators of cellular phosphate status. In human cells, 1,5-InsP_8_ binds to the SPX domain of XPR1, an interaction required for XPR1-mediated inorganic phosphate export ^31,34^. As a consequence of this feedback loop, defective phosphate export leads to accumulation of inositol pyrophosphates, as previously shown in HAP1 cells^36^ and likely occurring in the *XPR1*-mutated THP-1 cells.

**Figure 5.**
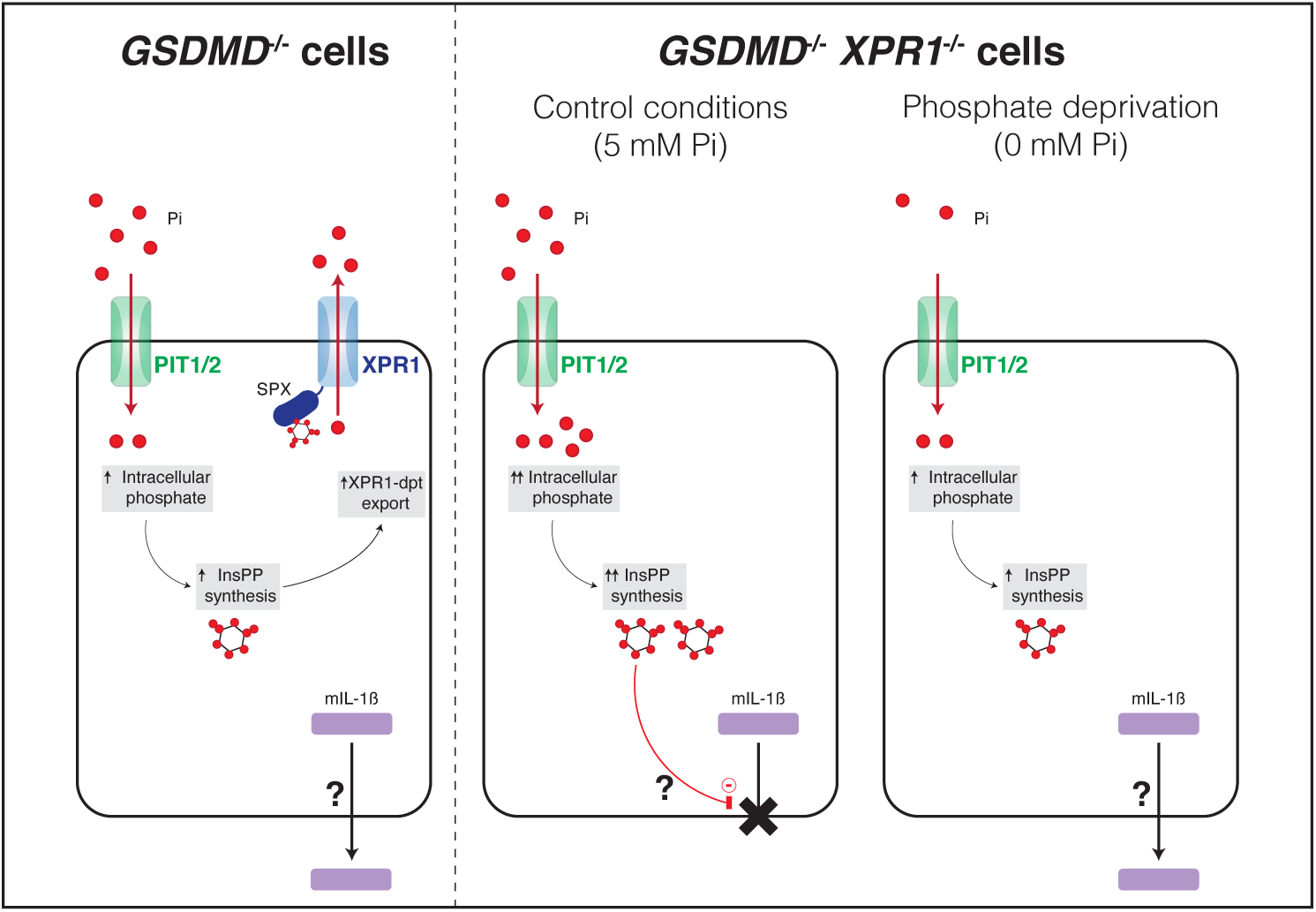
Proposed model linking phosphate homeostasis to mature IL-1β secretion. In *GSDMD*^-/-^ THP-1 cells phosphate homeostasis is regulated by the phosphate importers PiT1 and PiT2 together with the phosphate exporter XPR1. Increased extracellular inorganic phosphate (Pi) is associated with elevated intracellular phosphate levels, which may promote the synthesis of inositol pyrophosphates, which in turn activate XPR1-dependent phosphate export. In *GSDMD* ^-/-^ *XPR1* ^-/-^ THP-1 cells, phosphate export is impaired, leading to intracellular phosphate accumulation and potentially increased levels of inositol pyrophosphates. Under these conditions, the secretion of mature IL-1β is inhibited. Under phosphate-deprivation conditions, intracellular phosphate levels are expected to decrease, which may lower inositol pyrophosphates levels and is associated with an increase of mature IL-1β secretion.

Beyond phosphate regulation, inositol pyrophosphates regulate many cellular processes ^49^ including stress responses ^50,51^, autophagy ^52,53^, and vesicular trafficking ^54–57^. Notably, vesicular trafficking of autophagosomes and secretory lysosomes has also been proposed to contribute to the unconventional secretion of mature IL-1β ^58–60^.

Further work is needed to elucidate the mechanistic links between phosphate metabolism, inositol pyrophosphate and mature IL-1β secretion. Nonetheless, the model used in this study, THP-1 cells stimulated with LPS alone, provides a suitable framework for such future investigations.

Importantly, the identification of XPR1 as a regulator of mature IL-1β secretion reveals an unexplored avenue with broad implications for inflammatory processes and diseases.

## RESSOURCE AVAILABILITY

### Lead contact

Further information and requests for resources and reagents should be directed to and will be fulfilled by the lead contact, Tiphanie Gomard.

### Materials availability

This study did not generate new unique reagents.

### Data and code availability

RNAseq data are deposited at NCBI GEO with accession number GSE117162. (https://www.ncbi.nlm.nih.gov/geo/query/acc.cgi?acc=GSE117162). Any additional information required to reanalyze the data reported in this paper is available from the lead contact upon request. Raw data from figure 3 were deposited on Mendeley at https://data.mendeley.com/preview/3nj4b7xpjw?a=9675c319-50fa-4fe7-bab5-5929e2882b61.

## ACKNOWLEDGMENTS

The authors thank Nicolas Dauguet for flow cytometry experiments, Julie Stockis, Marc Hennequart and Laure Dumoutier for their critical reading of the manuscript and Suzanne Depelchin for expert administrative assistance. This work was funded by the Fonds National de la Recherche Scientifique-FNRS, Brussels, Belgium, and the WEL Research Institute, Belgium. C. S. was a FRIA grantee (FNRS).

## AUTHOR CONTRIBUTIONS

T.G. and P.C. conceived the project; N.R., C.S., S.T., E.V.S., JL.B., S.L., P.C. and T.G. designed the experiments; N.R., C.S. and T.G. performed the experiments and T.G and P.C. wrote the manuscript.

## SUPPLEMENTAL INFORMATION

Document S1. Figures S1-S7 and Table S1-S2

Datasets S1 and S2. Excel file containing additional data too large to fit in a PDF, related to Figure 3

## STAR★METHODS

### KEY RESOURCES TABLE

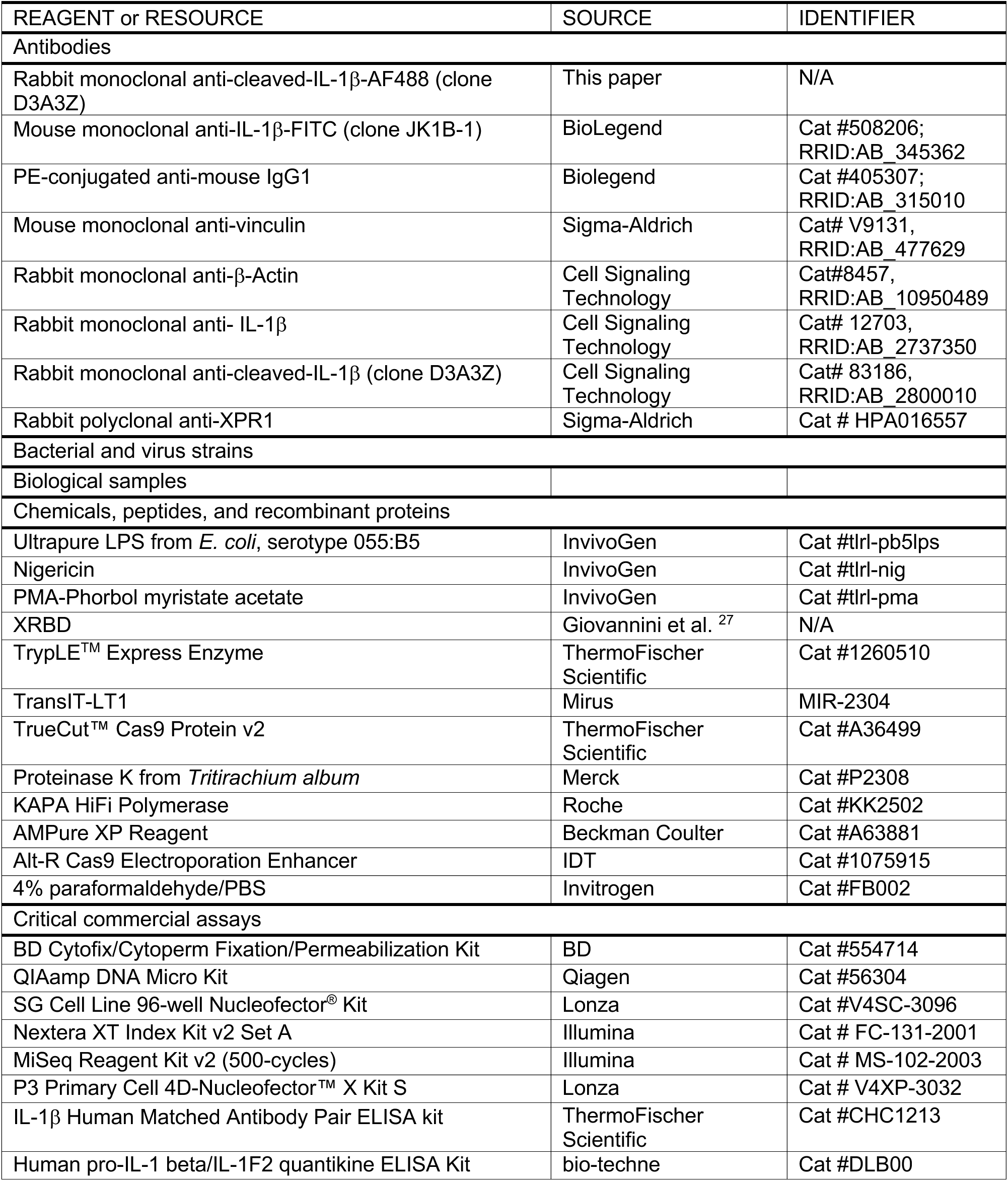
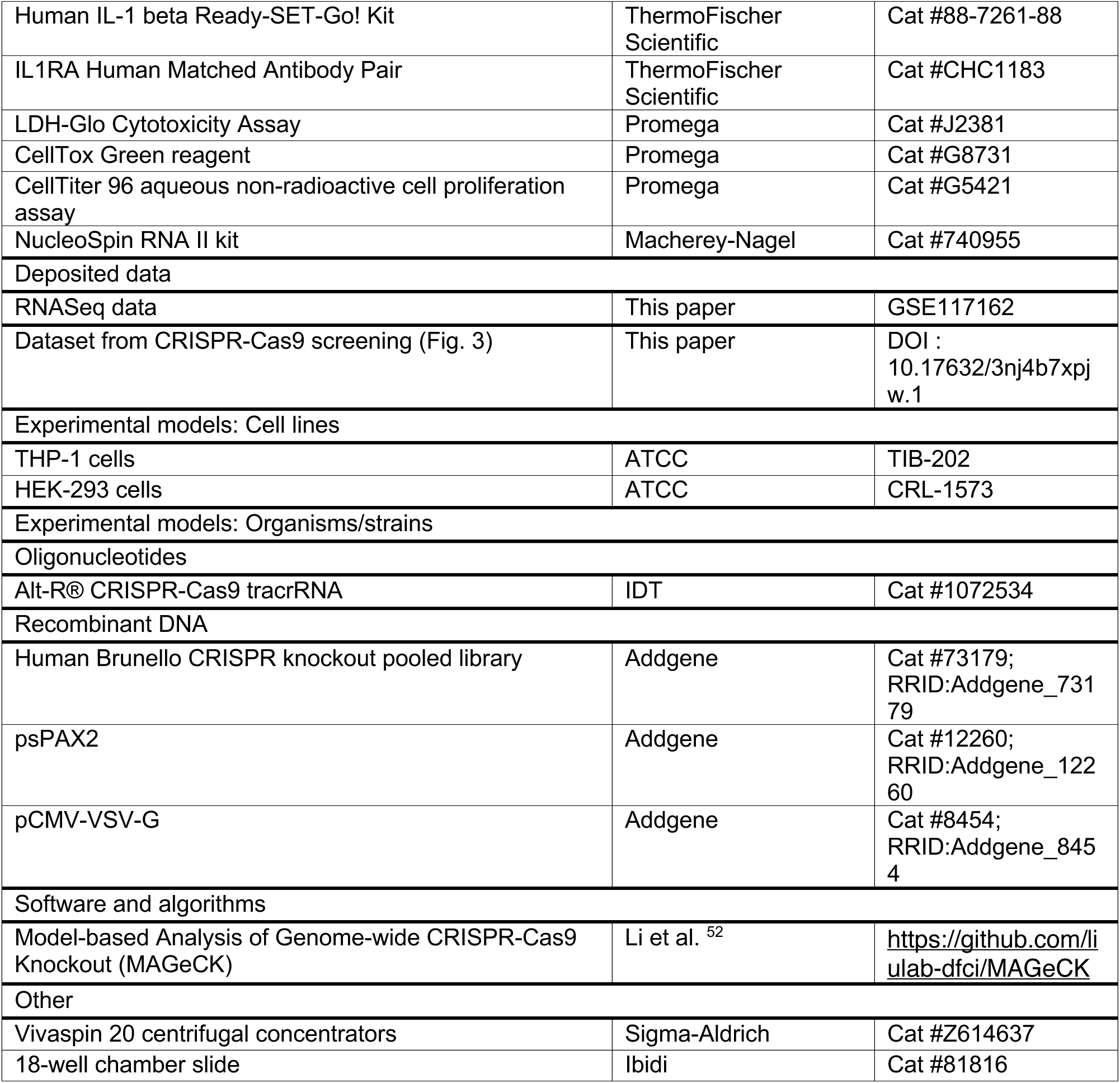

### METHOD DETAILS

#### Cell lines

THP-1 cells (ATCC TIB-202) were cultured in RPMI 1640 medium supplemented with L-glutamine, HEPES, 2-mercaptoethanol, 100 U/mL penicillin, 100 mg/mL streptomycin (Pen+Strep), and 10% (v/v) FBS. HEK293T cells were cultured in DMEM, low glucose supplemented with L-glutamine, sodium pyruvate, Pen+Strep, and 10% (v/v) FBS.

#### Reagents to be reconstituted

Ultrapure LPS from *E. coli*, serotype 055:B5, was purchased from InvivoGen (tlrl-pb5lps) and suspended in sterile endotoxin-free water at 5 mg/mL. Nigericin was purchased from InvivoGen (tlrl-nig) and resuspended in ethanol at 5 mg/mL (6.7 mM).

#### PMA-differentiation of THP-1 cells and cell stimulation

THP-1 cells were seeded at 10^5^ cells/200 µL per well in a flat-bottom 96 well TC microplate, differentiated for 3 h with 300 ng/mL PMA (InvivoGen, tlrl-pma), washed three times with warm test medium (phenol red-free RPMI 1640 supplemented with L-glutamine, 2-mercaptoethanol, Pen+Strep, and 5% (v/v) FBS), and incubated for 18 h in fresh test medium prior to LPS stimulation.

We observed that PMA treatment for 3 h followed by an 18 h rest period in the absence of PMA resulted in increased expression of *IL1B* and other genes involved in LPS response in THP-1 cells (Fig. S1). To limit background signals in LDH release and CellTox Green assays, cells were stimulated in test medium with 10 ng/mL ultrapure LPS from *E. coli* for 24 h. As a positive control for cell death, cells were treated with nigericin (10 µM) added 4 h after LPS.

For PMA differentiation and LPS stimulation under phosphate-deprivation conditions, the same protocol was used with custom phosphate-free RPMI 1640 medium (Gibco). Where indicated, test medium was supplemented with 5mM phosphate using sodium phosphate dibasic solution (Santa Cruz #sc-301828A; CAS Number 7558-79-4).

#### XRBD protein purification

We kindly thank Jean Luc Battini for providing the sequence for the XRBD-mFc construct published previously^29^. The plasmid encoding XRBD (strain NZB) was synthesized by ImmunoPrecise Antibodies. XRBD protein was expressed and purified by ImmunoPrecise Antibodies.

#### Flow cytometry

To select cells that accumulate mature IL-1β in the cytoplasm, cells were incubated for 10 min at 4 °C in PBS-2mM EDTA and scratched. Cells were fixed and permeabilized with the BD Cytofix/Cytoperm Fixation/Permeabilization Kit (BD #554714). Cells were resuspended in Perm Buffer, stained with coupled antibody against cleaved IL-1β (Cell Signaling – custom) at 0.3 µg/mL, and incubated overnight at 4 °C. Cells were washed two times in Perm Buffer and resuspended in PBS-2mM EDTA-1% FCS for cell sorting on a FACSAria III cell sorter (BD Biosciences).

We verified the specificity of the staining using HEK-293 cells transfected with plasmids encoding either human mature IL-1β or pro-IL-1β (Fig. S6). Transfected HEK293 cells were detached with TrypLE^TM^ Express Enzyme (ThermoFischer Scientific #1260510) for 10 min. Cells were fixed and permeabilized with the BD Cytofix/Cytoperm Fixation/Permeabilization Kit (BD #554714). Cells were resuspended in Perm Buffer, stained with different coupled antibody at 10 µg/mL, and incubated 30min at room temperature. We used a rabbit IgG Isotype Control (Alexa Fluor 488 conjugate) (Cell Signaling #4340) as negative control (Isoctrl), a rabbit Alexa Fluor 488 coupled antibody against cleaved IL-1β (Cell Signaling – custom) and a mouse FITC anti-human IL-1β antibody (BioLegend #508206, clone JK1B-1) which recognizes both pro- and mature IL-1β. Cells were washed two times in Perm Buffer and resuspended in PBS-2mM EDTA-1% FCS for analysis on a LSRFortessa flow cytometer (BD Biosciences).

Detection of XPR1 at the plasma membrane was performed as previously described^36^, using soluble RBD derived from X-MLV retrovirus and fused to a mouse IgG1 Fc domain (XRBD) and able to bind XPR1. XRBD was produced and used as described previously^29^. Briefly, 1 × 10^5^ cells were washed in PBS-2 mM EDTA-1% FCS, resuspended in 100 µL of PBS-2 mM EDTA-1% FCS containing 0.1 µg/mL of XRBD, and incubated at 37 °C for 30 min. Cells were then washed twice with cold PBS-2 mM EDTA-1% FCS and labeled for 20 min on ice with PE-conjugated anti-mouse IgG1 antibodies (2 µg/mL; BioLegend, #405307). Cells were then washed in PBS-2 mM EDTA-1% FCS and analyzed on a LSRFortessa flow cytometer (BD Biosciences). Data analysis was performed using FlowJo software.

#### Genome-wide CRISPR knockout screening

The human Brunello CRISPR knockout pooled library was a gift from David Root and John Doench (Addgene #73179)^61^. The gRNA pooled library in lentiCRISPRv2 was amplified and sequenced according to the accompanying online manual (https://www.addgene.org/pooled-library/broadgpp-human-knockout-brunello/). For production of the lentiviral pooled library, 60 × 10^6^ HEK293T cells were transfected with the plasmids psPAX2 (Addgene, #12260) and pCMV-VSV-G (Addgene, #8454) and the Brunello library in LentiCRISPRv2, in 10-cm dishes at a confluence of 70% with TransIT-LT1 (Mirus, MIR-2304) according to the provider’s protocol. Viral supernatants were collected 48 h post-transfection and passed through a 0.45-µm PVDF filter via a syringe, and viruses were concentrated 25-fold using Vivaspin 20 centrifugal concentrators (Sigma-Aldrich, Z614637). A total of 60 × 10^6^ THP-1 cells were infected with fresh viruses (at a multiplicity of infection of 0.9), with amplification for 19 days. A total of 3 × 50.10^6^ infected-THP-1 cells were differentiated with PMA and stimulated with LPS for 24 h (as described in the previous protocol). LPS-stimulated cells were stained with anti-cleaved-IL-1β-AF488–coupled antibody (as described previously), and positive IL-1β cells were sorted in triplicate (Fig. S4). Genomic DNA from sorted cells was isolated with the QIAamp DNA Micro Kit (Qiagen #56304) according to the manufacturer’s recommendation. As control, genomic DNA was isolated from 50 × 10^6^ cells before PMA differentiation and LPS stimulation by salting out. gRNA inserts were amplified via PCR using primers harboring Illumina Nextera XT adapter sequences, and the resulting libraries were sequenced on an Illumina MiSeq platform (platform CTMA, UCLouvain) using the MiSeq Reagent Kit v2, 500 cycles (Illumina). MaGeCK analysis ^62^ ^63^ of the sequencing count resulted in individual sgRNA scores (Dataset S1) as well as gene scores (Dataset S2) for significant enrichment within selected cells versus control cells. Raw data were deposited on Mendeley at https://data.mendeley.com/preview/3nj4b7xpjw?a=9675c319-50fa-4fe7-bab5-5929e2882b61.

#### Generation of knockout THP-1 cells

For genetic loss-of-function studies, a guide targeting *GSDMD*, *XPR1*, or *KIDINS220* (sequences in Table S1) was used for ctRNP complexes. Guide RNA complexes were formed by combining the crRNA and tracrRNA (IDT #1072534) in equal molar amounts in IDT Duplex Buffer at 100 µM by heating the oligos at 95 °C, followed by slow cooling to room temperature. The ctRNP complex was prepared by combining 300 pmol of the crRNA:tracrRNA complex with 60 pmol of Cas9 protein (TrueCut Cas9 Protein v2, ThermoFisher, #A36499), with an incubation for 15 min at room temperature. THP-1 cells were nucleofected with the complex according to instructions from Amaxa. Briefly, 10^6^ THP-1 cells were washed in PBS, resuspended in 20 µL of Nucleofection Solution SG (Lonza #V4SC-3096), mixed with 5 µL of the ctRNP complex, and nucleofected with the Amaxa 4D-Nucleofector (Lonza) with the protocol EN-138. We directly added 80 µL of warm medium and seeded cells in 1 mL in a 24-well plate. Cells were amplified and cloned one week later with the BD FACS Aria III at the Flow Cytometry and Cell Sorting Facility of the de Duve Institute, with one cell per well into 96-well plates. After 3 weeks, growing clones (22 per nucleofection) were amplified in M48 wells, and genomic DNA was extracted by boiling and digestion with Proteinase K at 10 µg/mL (Merck #P2308). Genomic DNA was amplified using PCR primers that flanked the crRNA cleavage site, generating an amplicon of approximatively 400 bp. Primers contained universal “tails” at the 5’ ends that allowed for a second amplification step to incorporate Illumina adapter sequences as well as sample-specific barcodes. PCR amplicons were generated using KAPA HiFi Polymerase (Roche #KK2502). Amplicons were purified by solid phase reversible immobilization bead cleanup using the 1.2x AMPure XP (Beckman Coulter #A63881) per the manufacturer’s protocol. NGS platform-specific adapters were incorporated in a subsequent amplification with universal primers (Nextera XT Index Kit v2, Illumina). After purification, the library was sequenced on an Illumina MiSeq platform (platform CTMA, UCLouvain) using the MiSeq Reagent Kit v2, 500 cycles (Illumina). Gene editing was guided by bioinformatics analysis through alignment of paired-end reads and programmatic evaluation of variants relative to the WT sequence. For each cell, we generated two different clones obtained with different gRNAs (Table S1), and for each clone, complete gene invalidation was verified by sequencing (Table S1).

#### Generation of mutated XPR1 THP-1 cells by CRISPR-mediated homology-directed repair

To generate mutated XPR1 THP-1 cells, we followed the same protocol as for the generation of knockout cells with some adjustments. A total of 10^6^ THP-1 cells were washed in PBS, resuspended in 20 µL of Nucleofection Solution P3 (Lonza #V4XP-3032), mixed with 5 µL of the ctRNP complex + 1.2 µL Alt-R Cas9 Electroporation Enhancer (IDT #1075915) + 1.2 µL HDR Donor oligo (100 µM), and nucleofected with the Amaxa 4D-Nucleofector (Lonza) with the protocol FF18. After nucleofection, cells were seeded in medium supplemented with HDR Enhancer V2 (IDT #10007910) for 24 h. For each clone, complete gene invalidation or mutation was verified by sequencing (Table S2).

#### ELISA

Supernatants were collected at the indicated times in non–TC-treated 96-well plates and centrifuged to eliminate residual cells. Supernatants were then transferred to a new plate and kept at -20 °C. To compare the amounts of secreted and intracellular mature IL-1β, after the collection of culture supernatants, cells were resuspended in 200 µL of medium (same volume as supernatants) and lysed by one cycle of freeze/thaw at -20 °C. Human mature IL-1β was quantified in supernatants and inside cells with the IL-1β Human Matched Antibody Pair ELISA kit from ThermoFisher Scientific (#CHC1213), according to the provider’s protocol. We verified that this kit recognizes mature human IL-1β and that pro-IL-1β is not detected at all (Fig. S7).

HEK293T cells were transfected with plasmids encoding either human mature IL-1β or pro-IL-1β. After 3 days, cells were either collected in the Laemmli buffer (reducing conditions) for western blot analysis or lysed by one cycle of freezing/thawing. These thawed cell lysates were centrifuged and the supernatant collected for ELISA. We measured pro-IL-1β with the Human pro-IL-1 beta/IL-1F2 quantikine ELISA Kit (bio-techne #DLB00) and human IL-1β (mature and pro-IL-1β) with the Human IL-1 beta Ready-SET-Go! Kit (ThermoFisher Scientific #88-7261-88), according to the provider’s protocol.

Human IL-1RA was quantified in supernatants and inside cells with the IL1RA Human Matched Antibody Pair from ThermoFisher Scientific (#CHC1183), according to the provider’s protocol.

#### Immunoblotting

Total cell lysates were prepared using Laemmli buffer containing β-mercaptoethanol, denaturated for 10 min at 95°C, separated by Bis-Tris denaturing SDS-PAGE and transferred onto nitrocellulose membranes (ThermoFisher Scientific). Membranes were blocked with 5% non-fat dry milk in PBS with 0,1% Tween 20 (PBST) for 1h at RT, and incubated overnight at 4°C with indicated primary antibodies diluted in 5% BSA/PBST at 1:1000 for anti-Vinculin (Sigma-Aldrich, #V9131), anti-IL-1β (Cell Signaling #12703), and anti-cleaved human IL-1β (Cell Signaling #83186). After washes, membranes were incubated 1h at RT with anti-mouse-HRP (R&D Systems, #HAF007) or anti-rabbit-HRP (Cell Signaling #7074). Chemoluminescent signals were recorded with a CCD-camera (Fusion Solo S, Vilber).

#### LDH release assay

A total of 15 µL of each supernatant was diluted in 135 µL of storage buffer (200 mM Tris-HCl, pH 7.3), 10% glycerol, and 1% BSA, according to the manufacturer’s instructions). Residual supernatant was harvested, and cells were resuspended in 200 µL medium supplemented with 1% Triton X-100 and incubated for 15 min at room temperature to obtain maximal release of cellular LDH. A total of 15 µL of lysed cells were diluted in 135 µL of storage buffer. Diluted supernatants and cells were kept at -20 °C. LDH activity was measured with the LDH-Glo Cytotoxicity Assay (Promega, #J2381) according to the manufacturer’s instructions.

Luminescence was measured using the Glomax Discover device. The percentage of LDH release was 100 × (Qty LDH in supernatants (mU/mL)) / (Qty LDH in supernatants (mU/mL) + Qty LDH in cells (mU/mL)).

#### CellTox^TM^ Green uptake assay

Cells were seeded in a flat-bottom 96-well TC white microplate, differentiated with PMA, and stimulated with LPS (as described in the previous protocol). Concentrated CellTox Green reagent was added to cells to a final dilution of 1:1000 (as recommended, Promega #G8731).

Fluorescence was measured at any point between 0 and 24 h using the Glomax Discover device, and the plate was returned to the incubator between reads. After 24 h, cells were lysed using Triton X-100 (0.2% final) and frozen. Recorded fluorescence after cell lysis represented the maximal fluorescence. The percentage of CTG uptake was 100 × (fluorescence at time T) / (maximal fluorescence).

#### Cell viability assay

Cell viability was measured with the CellTiter 96 aqueous non-radioactive cell proliferation assay from Promega (#G5421), according to the manufacturer’s protocol. At the indicated times, supernatants were discarded and replaced by 100 µL of warm medium plus 20 µL of MTS/PMS mix. After a 2-h incubation at 37 °C under 5% CO_2_, light absorbance was recorded at 490 nm.

#### Immunofluorescence

Cells were seeded in an 18-well chamber slide for microscopy (10^5^ cells/100 µL, Ibidi #81816), differentiated with PMA, and stimulated with LPS (as described). After 6 h of LPS stimulation, cells were washed with PBS, fixed for 15 min in 4% paraformaldehyde/PBS (Invitrogen #FB002), washed with PBS, incubated for 15 min in permeabilization buffer (PBS-0.5% Triton X100, Sigma X-100), and incubated for 30 min in blocking buffer (PBS-1% BSA-1% goat serum-0.1% Triton X-100; BSA, Sigma #A7030; goat serum, Millipore S26). Cells were then incubated overnight at 4 °C in the first antibody diluted 1:100 in blocking buffer (anti-XPR1 Sigma Prestige #HPA016557), washed 3 × 5 min in PBS-0.1% Triton and incubated for 1 h at room temperature with an Alexa 488–conjugated secondary antibody (Invitrogen #A48282). All images were acquired at the Platform for Imaging Cells and Tissues of the de Duve Institute (UCLouvain-Brussels) with the point scanner confocal and multiphoton microscope (Zeiss LSM980) with a 63× objective. Images were processed using the ImageJ software-based Fiji platform.

#### Library preparation for RNAseq

RNA was isolated using the NucleoSpin RNA II kit according to manufacturer’s instructions (Macherey-Nagel, #740955). Libraries were prepared by Integragen Genomics with the TruSeq Stranded mRNA kit protocol according to the supplier’s recommendations. Briefly, we performed purification of poly-A containing mRNA molecules using poly-T–coated magnetic beads from 1 µg of total RNA, with fragmentation using divalent cations under elevated temperature to obtain approximately 300-bp fragments; double-stranded cDNA synthesis; Illumina adapter ligation; cDNA library PCR amplification; and 100-bp paired-end sequencing on an Illumina HiSeq4000.

#### RNAseq analysis

Alignment to the GRCh38 reference genome was done using STAR. Bam files were sorted with Samtools 1.6. Gene counts were obtained using HTSeq 0.9.1 htseq-count (union mode). Count normalization was performed with DESeq2_1.18.1. The RNAseq data are deposited at NCBI GEO with accession number GSE117162 (https://www.ncbi.nlm.nih.gov/geo/query/acc.cgi?acc=GSE117162).

