## Supplemental file for "The phosphate exporter XPR1 regulates a gasdermin D–independent mature IL-1β secretion pathway in LPS-stimulated human monocytic cells"

**Fig. S1. Related to Figure 1, Transcriptomic analysis of THP-1 cells following differentiation with PMA.**

THP-1 cells were treated or not with PMA (300 ng/mL) for 3 h, washed, and further differentiated for 18 h prior to RNAseq analysis. Gene expression levels (read counts) in PMA-treated cells are indicated on the right.

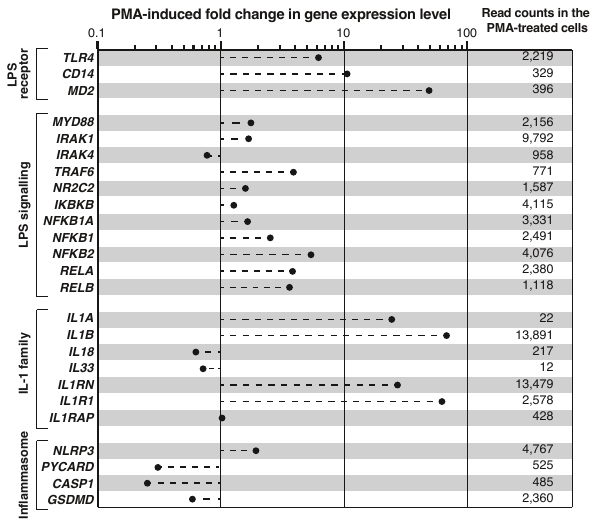

**Fig. S2. Related to Figure 1, Pro-IL-1β processing in THP-1 stimulated with LPS alone is dependent on both caspase 1 and NLRP3, and independent of caspase 8, 4 or 5.**

PMA-differentiated THP-1 cells (WT or *CASP1*, *NLRP3*, *CASP8*, *CASP4* or *CASP5* KO) were stimulated with LPS (10 ng/mL) or not for 24 h, nigericin (10µM) was added 4 h after LPS. At the end of the stimulation, supernatants were collected, and cells were resuspended in 200 µL of medium and lysed by single freeze-thaw cycle. Samples for immunoblotting were prepared using LAEMMLI Buffer (4x) containing β-mercaptoethanol, denaturated for 10 min at 95°C, separated by Bis-Tris denaturing SDS-PAGE and transferred onto nitrocellulose membranes.

Top panel. Detection of proIL-1β and mature IL-1β in cell lysates and corresponding culture supernatants. Membranes were probed with the indicated antibodies.

Immunoblots were intentionally overexposed to demonstrate the absence of mature IL-1β in *CASP1*^-/-^ and *NLRP3*^-/-^ THP-1 cells. The presence of proIL-1β and mature IL-1β in unstimulated cells was clone-dependent. As shown in Figure S1, PMA-differentiation upregulated pro-IL-1β expression.

Bottom panel. Concentrations (ng/mL) of mature IL-1β measured in the supernatants of the corresponding samples by ELISA.

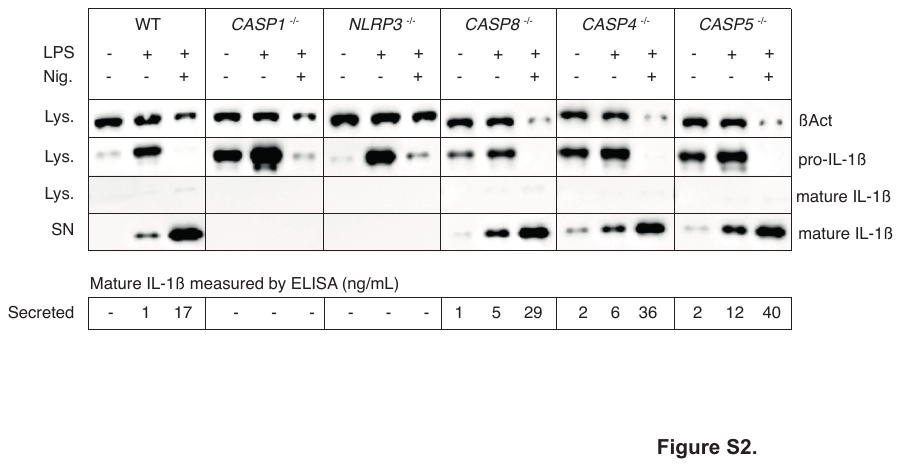

**Fig. S3. Related to Figure 2, Immunoblot confrmation of mature IL-1β accumulation in GSDMD^-/-^ THP-1 cells after LPS stimulation.**

PMA-differentiated THP-1 cells (WT or *GSDMD^-/-^*) were stimulated with LPS (10 ng/mL) or not for 24 h, nigericin (10µM) was added 4 h after LPS. Cells were lysed at 5 h and 24 h in recuing Laemmli buffer and analyzed by Western blotting.

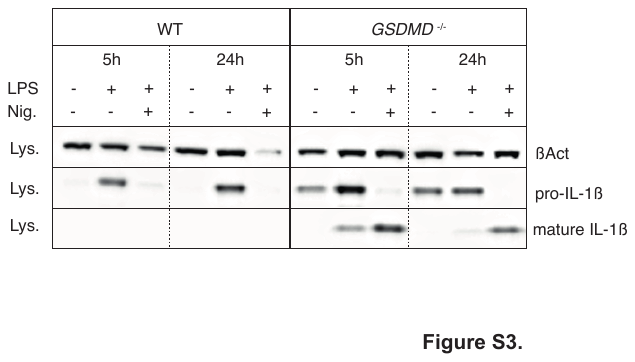

Fig. S4. Related to Figure 3, Mature IL-1β–enriched cell sorting and quality control of the CRISPR/Cas9 screening.

(A) Results of cell sorting performed during the CRISPR/Cas9 screening. For visualization purposes, replicates 1, 2, and 3 were downsampled to 30,000 cells (total number of analyzed cells: Rep1 – 21,764,670; Rep2 – 19,624,963; Rep3 – 19,918,967).

(B) Quality control analysis using MAGeCK. Percentages of reads mapped to the Brunello library were >65% (consistent with standard for this type of screen). Library representation (defined as the proportion of different gRNAs from the library detected among the mapped reads) was close to 100% in control cell populations.

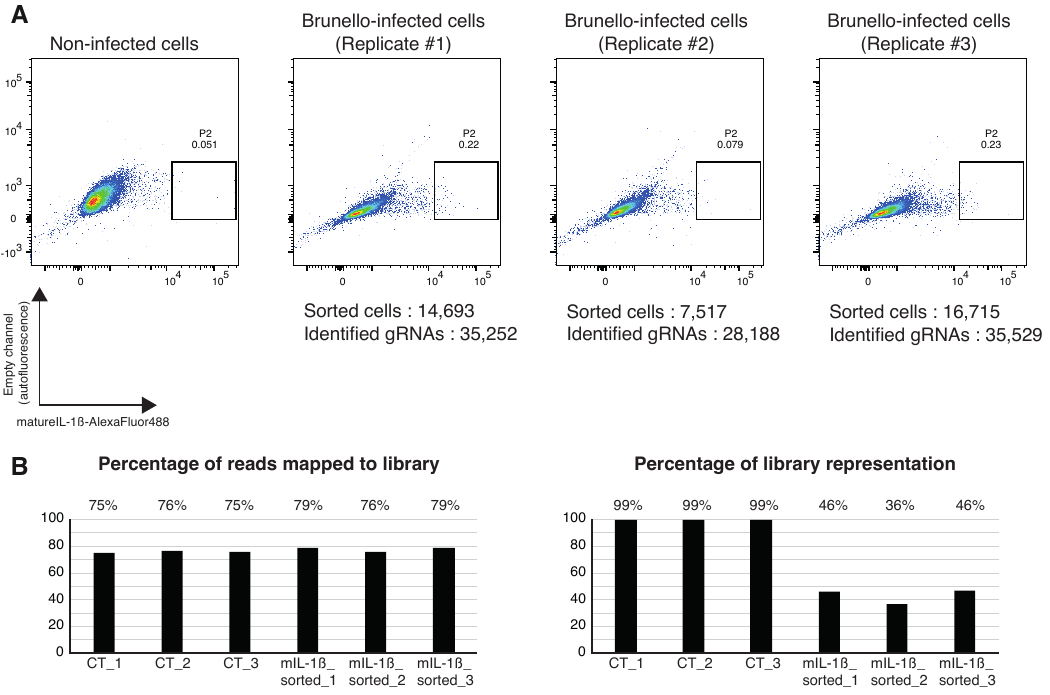

Fig. S5. Related to Figure 4, XPR1 positively regulates GSDMD-independent mature IL-1β secretion efficiency.

(A) Raw data used to calculate IL-1β secretion efficiencies. For each replicate of each independent experiment, the amounts of mature IL-1β (ng/10^5^ cells) measured in supernatants (SN) and within cells are indicated, together with the corresponding calculated secretion efficiencies (Eff). Only secretion efficiency values are shown in Fig. 4.

(B) Representative immunofluorescence images of *GSDMD*^-/-^ *KIDINS220*^-/-^, *GSDMD*^-/-^ *XPR1^L218S/L218S^*, and *GSDMD*^-/-^ *XPR1^KSC/KSC^* THP-1 cells differentiated with PMA and stained with anti-XPR1 (green) and co-labeled with DAPI (blue). Magnification 63×.

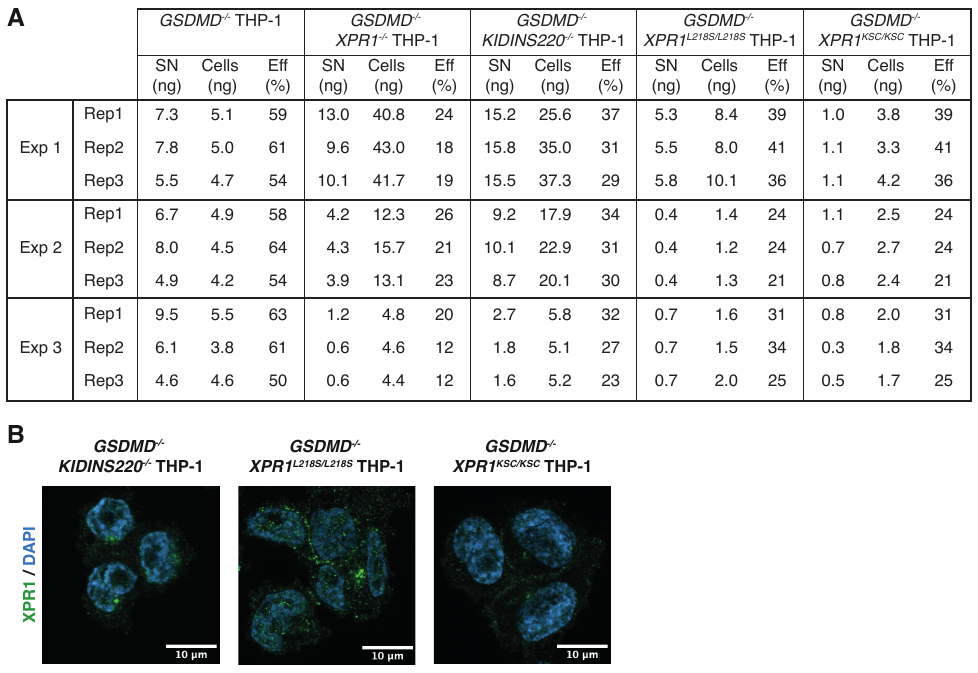

Fig. S6. Validation that the antibody used for the CRISPR library screening specifically recognizes human mature IL-1β but not pro-IL-1β.

HEK293T cells were transfected with plasmids encoding either human mature IL-1β or pro-IL-1β. After 3 days, cells were collected, fixed, permeabilized, and stained with a rabbit IgG Isotype Control (Alexa Fluor 488 (AF488) conjugate) as negative control (Isoctrl), a rabbit anti-cleaved IL-1β antibody conjugated to Alexa Fluor 488 (AF488) or a mouse FITC-conjugated anti-human IL-1β antibody recognizing both pro- and mature IL-1β. The AF488-conjugated anti-cleaved IL-1β specifically detected mature IL-1β into cells.

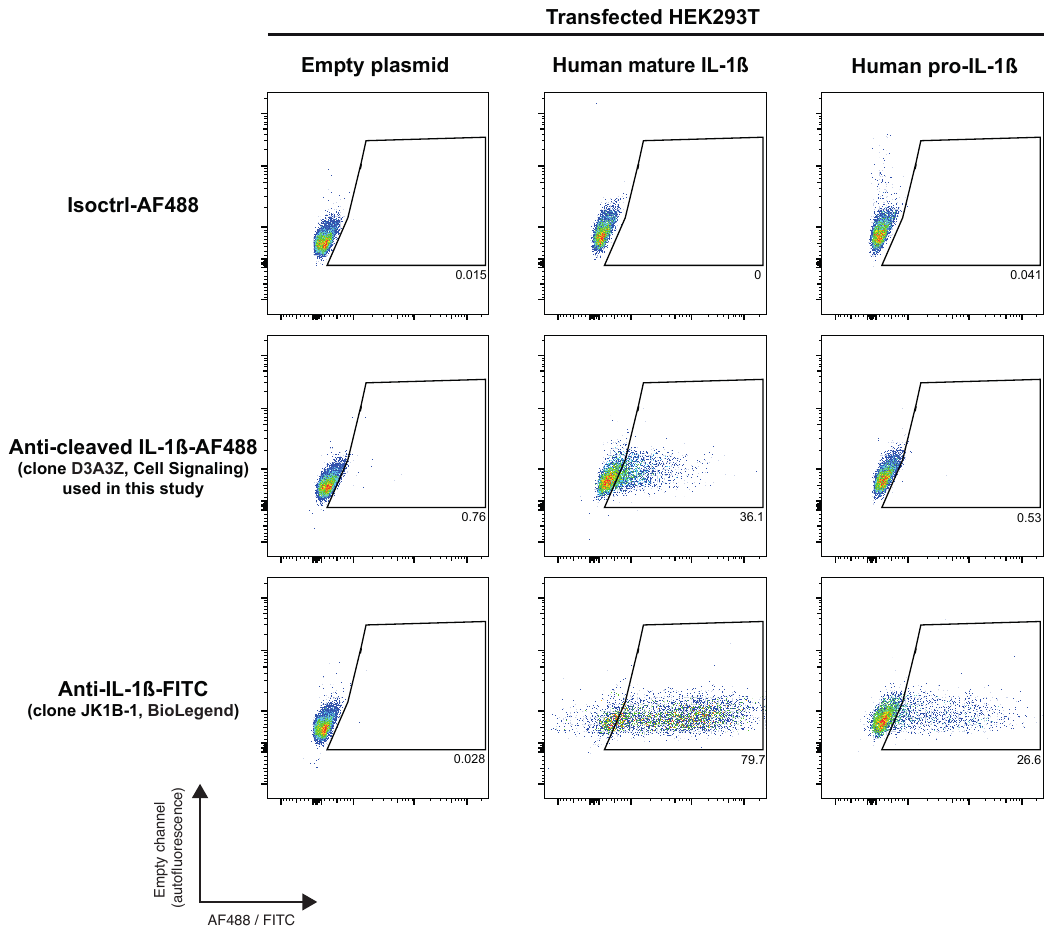

Fig. S7. Validation that the human IL-1β Cyto-Set ELISA kit detects human mature IL-1β but not pro-IL-1β.

HEK293T cells were transfected with plasmids encoding either human mature IL-1β or pro-IL-1β. After 3 days, cells were either collected in the Laemmli buffer (reducing conditions) for western blot analysis or lysed by a single freeze-thaw cycle. These thawed cell lysates were centrifuged and the supernatant collected for ELISA.

Top panels. Detection of pro-IL-1β and mature IL-1β in transfected HEK293T cells by immunoblotting. A single membrane was cut into three pieces and probed with the indicated antibodies.

Bottom panel. Concentrations (pg/mL) of IL-1β measured in cell lysates with three different ELISA kits.

The Quantikine pro-IL-1β kit detected pro-IL-1β but not mature IL-1β. The Ready-SET-Go! kit detected both mature IL-1β and pro-IL-1β. The Human IL-1β Cyto-Set kit detected mature IL-1β and produced only a very weak signal in lysates from HEK293T cells expressing pro-IL-1β. We cannot exclude the possibility that this signal reflects a low level of pro-IL-1β processing in HEK293T cells. The Human IL-1β Cyto-Set kit was used throughout this study.

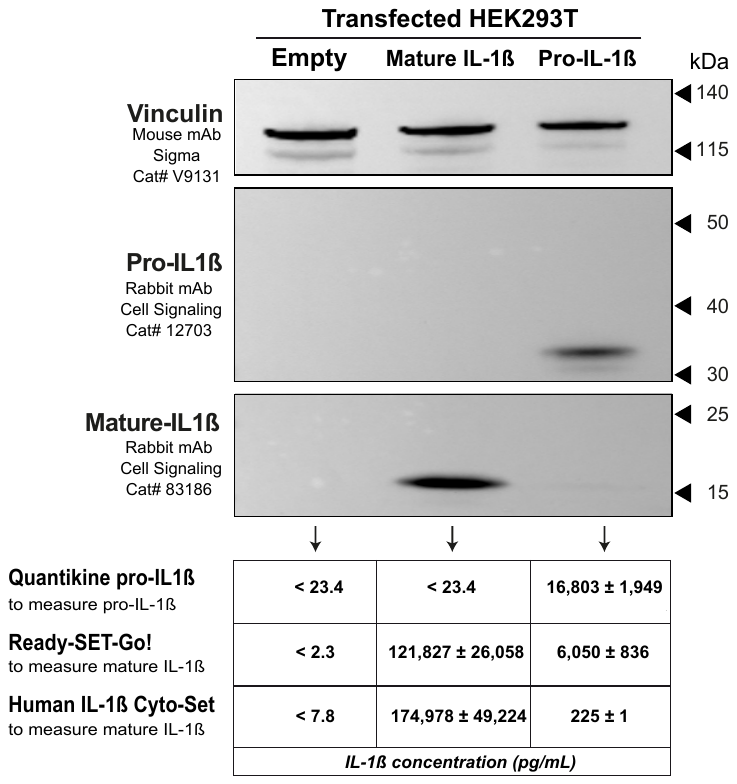

Table S1. Genetic validation by targeted sequencing of all gene-invalidated THP-1 cell clones used in this study.

For each clone, the WT reference sequence of the targeted gene is shown on top with the sgRNA target site indicated in red. Sequences of the edited alleles are shown below; deletions are represented by dots, and insertions are indicated in a larger font.

(1) Clone used for data shown in Figs. 1–4.

(2) Clone used for data shown in Figs. 3 and 4.

(3) Clone used for data shown in Fig. 4.

(4-8) Clones used for data shown in Fig. S2.

| **THP-1 clone** | **Gene** | **Sequence** | | **Length of encoded protein (AA)** |
| --- | --- | --- | --- | --- |
| ***GSDMD*^-/-^ cl1 ^(1)^** | *GSDMD* | WT | ATGGGGTCGGCCTTTGAGCGGGTAGTCCGGAGAGTGGTCCAGGAGCTGGACCAT | 484 |
|  |  | Allele 1 | ATGGGGTCGGCCTTTG**A**AGCGGGTAGTCCGGAGAGTGGTCCAGGAGCTGGACCAT | 41 |
|  |  | Allele 2 | ATGGGGTCGGCCTTTGA**..**GGGTAGTCCGGAGAGTGGTCCAGGAGCTGGACCAT | 40 |
| *GSDMD*^-/-^ cl2 | *GSDMD* | WT | CAGGAGCTGGACCATGGTGGGGAGTTCATCCCTGTGACCAGCCTGCAGAGCTCCACT | 484 |
|  |  | Allele 1 | CAGGAGCTGGACCATGG**.**GGGGAGTTCATCCCTGTGACCAGCCTGCAGAGCTCCACT | 24 |
|  |  | Allele 2 | CAGGAGCTGGACCATG**GT**GTGGGGAGTTCATCCCTGTGACCAGCCTGCAGAGCTCCACT | 25 |
| GSDMD^-/-^ XPR1^-/-^ cl1 | *XPR1* | WT | TACTAGATTGCTGGATTCCTCGGGATATTGTGGTGCCTGAGCCTTCTGGCATGCTTCTTT | 696 |
|  |  | Allele 1 | TACTAGATTGCTGGATTCC**........**TTGTGGTGCCTGAGCCTTCTGGCATGCTTCTTT | 335 |
|  |  | Allele 2 | TACTAGATTGCTGGATTCCTCGG**..**TATTGTGGTGCCTGAGCCTTCTGGCATGCTTCTTT | 337 |
|  |  | Allele 3 | TACTAGATTGCTGGATTCCTCGGG**.**TATTGTGGTGCCTGAGCCTTCTGGCATGCTTCTTT | 327 |
|  | *GSDMD* | WT | ATGGGGTCGGCCTTTGAGCGGGTAGTCCGGAGAGTGGTCCAGGAGCTGGACCAT | 484 |
|  |  | Allele 1 | ATGGGGTCGGCCTTT**.**AGCGGGTAGTCCGGAGAGTGGTCCAGGAGCTGGACCAT | 7 |
|  |  | Allele 2 | ATGGGGTCGGCCTTTGA**A**GCGGGTAGTCCGGAGAGTGGTCCAGGAGCTGGACCAT | 41 |
|  |  | Allele 3 | ATGGGGTCGGCCTTT**.....**GGTAGTCCGGAGAGTGGTCCAGGAGCTGGACCAT | 39 |
| **GSDMD^-/-^ XPR1^-/-^ cl2 ^(2)^** | *XPR1* | WT | GAAGTTCGCCGAGCACCTCTCCGCGCACATCACTCCCGAGTGGAGGAAGCAATACATCCA | 696 |
|  |  | Allele 1 | GAAGTTCGCCGAGCACCTCTCCGCGC**....................**AAGCAATACATCCA | 15 |
|  |  | Allele 2 | GAAGTTCGCCGAGCACCTCTCCGCGCACATCACT**.**CCGAGTGGAGGAAGCAATACATCCA | 46 |
|  | *GSDMD* | WT | ATGGGGTCGGCCTTTGAGCGGGTAGTCCGGAGAGTGGTCCAGGAGCTGGACCAT | 484 |
|  |  | Allele 1 | ATGGGGTCGGCCTTT**..**GCGGGTAGTCCGGAGAGTGGTCCAGGAGCTGGACCAT | 40 |
|  |  | Allele 2 | ATGGGGTCGGCCTTTG**.**GCGGGTAGTCCGGAGAGTGGTCCAGGAGCTGGACCAT | 7 |
|  |  | Allele 3 | ATGGGGTCGGCCTTTGA**..**GGGTAGTCCGGAGAGTGGTCCAGGAGCTGGACCAT | 40 |
| **GSDMD^-/-^ KIDINS220^-/-^ cl1 ^(3)^** | *KIDINS220* | WT | TGTAGTGGGATTGGCCTTTGTGTTGAACTGTCGTACATGGTGGCAAGTGCTGGACTCGCT | 1771 |
|  |  | Allele 1 | TGTAGTGGGATTGGCCTTTGTGTTGAACTGTCGTAC**.**TGGTGGCAAGTGCTGGACTCGCT | 718 |
|  |  | Allele 2 | TGTAGTGGGATTGGCCTTTGTGTTGAACTGTCGTACA**.**GGTGGCAAGTGCTGGACTCGCT | 718 |
|  | *GSDMD* | WT | ATGGGGTCGGCCTTTGAGCGGGTAGTCCGGAGAGTGGTCCAGGAGCTGGACCAT | 484 |
|  |  | Allele 1 | ATGGGGTCGGCCTTTGA**.**CGGGTAGTCCGGAGAGTGGTCCAGGAGCTGGACCAT | 7 |
|  |  | Allele 2 | ATG**................**GGGTAGTCCGGAGAGTGGTCCAGGAGCTGGACCAT | 2 |
|  |  | Allele 3 | ATGGGGTCGGCCTTTG**A**AGCGGGTAGTCCGGAGAGTGGTCCAGGAGCTGGACCAT | 41 |
| GSDMD^-/-^ KIDINS220^-/-^ cl2 | *KIDINS220* | WT | GTGCACATCGTAGAGGAACTACTGAAATGTGGGGTTAACTTGGAGCACCGTGATATGGGAGGA | 1771 |
|  |  | Allele 1 | GTGCACATCGTAGAGGAACTACTGAAATGTGG**.....**ACTTGGAGCACCGTGATATGGGAGGA | 98 |
|  |  | Allele 2 | GTGCACATCGTAGAGGAACTACTGAAATGTGGGGTT**A**AACTTGGAGCACCGTGATATGGGAGGA | 100 |
|  | *GSDMD* | WT | ATGGGGTCGGCCTTTGAGCGGGTAGTCCGGAGAGTGGTCCAGGAGCTGGACCAT | 484 |
|  |  | Allele 1 | ATGGGGTCGGCCTTTG**.**GCGGGTAGTCCGGAGAGTGGTCCAGGAGCTGGACCAT | 7 |
|  |  | Allele 2 | ATGGGGTCGGCCTTTGA**.....**TAGTCCGGAGAGTGGTCCAGGAGCTGGACCAT | 39 |
| **CASP1 ^-/-^ cl1 ^(4)^** | *CASP1* | WT | AGGTCCTGAAGGAGAAGAGAAAGCTGTTTATCCGTTCCATGGGTGAAGGTACAATAAATG | 404 |
|  |  | Allele 1 | AGGTCCTGAAGGAGAAGAGAAAGCTGTTTATCCGT**.......**GTGAAGGTACAATAAATG | 19 |
|  |  | Allele 2 | AGGTCCTGAAGGAGAAGAGAAAGCTGTTTATCC**.......**GGGTGAAGGTACAATAAATG | 19 |
| **CASP8 ^-/-^ cl1 ^(5)^** | *CASP8* | WT | GGCCTCCCTCAAGTTCCTGAGCCTGGACTACATTCCGCAAAGGAAGCAAGAACCCATCAA | 479 |
|  |  | Allele 1 | GGCCTCCCTCAAGTTCCTGAGCCTGGA**.......**CCGCAAAGGAAGCAAGAACCCATCAA | 40 |
|  |  | Allele 2 | GGCCTCCCTCAAGTTCCTGAGCCTGGACTA**.......**CAAAGGAAGCAAGAACCCATCAA | 40 |
|  |  | Allele 3 | GGCCTCCCTCAAGTTCCTGAGCCTGGACTACATTCCG**.**AAAGGAAGCAAGAACCCATCAA | 42 |
| **CASP4 ^-/-^ cl1 ^(6)^** | *CASP4* | WT | GGGTCATGGCAGACTCTATGCAAGAGAAGCAACGTATGGCAGGACAAATGCTTCTTCAAA | 377 |
|  |  | Allele 1 | GGGTCATGGCAGACTCTATGCAAGAGAAGCAACGT**.....**AGGACAAATGCTTCTTCAAA | 76 |
|  |  | Allele 2 | GGGTCATGGCAGACTCTATGCAAGAGAAGCAAC**.....**G**A**AGGACAAATGCTTCTTCAAA | 76 |
|  |  | Allele 3 | GGGTCATGGCAGACTC**........................................**CAAA | 66 |
| **CASP5 ^-/-^ cl1 ^(7)^** | *CASP5* | WT | AAAGATCACCAGTGTAAAACCTCTTCTGCAAATCGAGGCTGGACCACCTGAGTCAGCAGAATCTACAAATATACTCAAACTT | 434 |
|  |  | Allele 1 | AAAGATCACCAGTGTAAAACCTCTTCT**.....................................**ACAAATATACTCAAACTT | 484 |
| **NLRP3 ^-/-^ cl1 ^(8)^** | *NLRP3* | WT | AGACCATGTGGATCTAGCCACGCTAATGATCGACTTCAATGGGGAGGAGAAGGCGTGGGC | 1037 |
|  |  | Allele 1 | AGACCATGTGGATCTAGCC**.............**ACTTCAATGGGGAGGAGAAGGCGTGGGC | 101 |
|  |  | Allele 2 | AGACCATGT**...............................**GGGGAGGAGAAGGCGTGGGC | 95 |
|  |  | Allele 3 | AGACCATGTGGATCTAGCCACGC**T**TAATGATCGACTTCAATGGGGAGGAGAAGGCGTGGGC | 84 |

Table S2. Genetic validation of the GSDMD^-/-^ THP-1 clones knocked-in with XPR1 mutants.

For each clone, the WT reference sequences of the targeted genes are shown on top with the sgRNA target site indicated in red. For the *XPR1* L218S and KSC mutations, the HDR donor templates are shown, with the introduced mutation(s) in a larger font. For each of the four THP-1 clones shown, all sequencing reads spanning the targeted *XPR1* locus were identical and contained the desired mutation(s).

For each clone, the WT reference sequence of the targeted gene is shown on top with the sgRNA target site indicated in red. Sequences of the edited alleles are shown below; deletions are represented by dots, and insertions are indicated in a larger font.

(1) Clones used for data shown in Fig. 4.

| **THP-1 clone** | **Gene** | **Sequence** | | **Length of encoded protein (AA)** |
| --- | --- | --- | --- | --- |
| ***GSDMD^-/-^ XPR1^L218S/L218S^* cl1 *^(1)^*** | *XPR1* | WT | GAACTTGAAGATGGTGACAGACAAAAGGCTATGAAGCGTTTACGGGTCCCCCCTCTGGGAGCTGCTCAGGTTAGTATTTGGTTCCAGTAGTT | 696 |
|  |  | HDR donor template | GAACTTGAAGATGGTGACAGACAAAAGGCTATGAAGCGTT**C**ACGGGTCCCCCCTCTGGGAGCTGCTCAGGTTAGTATTTGGTTCCAGTAGTT | 696 |
|  | *GSDMD* | WT | ATGGGGTCGGCCTTTGAGCGGGTAGTCCGGAGAGTGGTCCAGGAGCTGGACCAT | 484 |
|  |  | Allele 1 | **..................**CGGGTAGTCCGGAGAGTGGTCCAGGAGCTGGACCAT | / |
|  |  | Allele 2 | ATGGGGTCGGCCTTTGA**A**GCGGGTAGTCCGGAGAGTGGTCCAGGAGCTGGACCAT | 40 |
| *GSDMD^-/-^ XPR1^L218S/L218S^* cl2 | *XPR1* | WT | GAACTTGAAGATGGTGACAGACAAAAGGCTATGAAGCGTTTACGGGTCCCCCCTCTGGGAGCTGCTCAGGTTAGTATTTGGTTCCAGTAGTT | 696 |
|  |  | HDR donor template | GAACTTGAAGATGGTGACAGACAAAAGGCTATGAAGCGTT**C**ACGGGTCCCCCCTCTGGGAGCTGCTCAGGTTAGTATTTGGTTCCAGTAGTT | 696 |
|  | *GSDMD* | WT | ATGGGGTCGGCCTTTGAGCGGGTAGTCCGGAGAGTGGTCCAGGAGCTGGACCAT | 484 |
|  |  | Allele 1 | ATGGGGTCGGCCTTTGA**.**CGGGTAGTCCGGAGAGTGGTCCAGGAGCTGGACCAT | 6 |
|  |  | Allele 2 | ATGGGGTCGGCCTTTGA**A**GCGGGTAGTCCGGAGAGTGGTCCAGGAGCTGGACCAT | 40 |
| ***GSDMD^-/-^ XPR1^KSC/KSC^* cl1 ^(1)^** | *XPR1* | WT | TCATTTCTTTCTGTAGAATCTGAATTTTACAGGGTTTCGAAAAATCCTGAAAAAGCATGACAAGATCCTGGAAACATCTCGTGGAGCAGATTGGCGAGTGGCTCACGTAGAG | 696 |
|  |  | HDR donor template | TCATTTCTTTCTGTAGAATCTGAATTTTACAGGGTTTCGA**GC**AATCCTG**GC**AAAGCATGAC**GC**GATCCTGGAAACATCTCGTGGAGCAGATTGGCGAGTGGCTCACGTAGAG | 696 |
|  | *GSDMD* | WT | ATGGGGTCGGCCTTTGAGCGGGTAGTCCGGAGAGTGGTCCAGGAGCTGGACCAT | 484 |
|  |  | Allele 1 | **..................**CGGGTAGTCCGGAGAGTGGTCCAGGAGCTGGACCAT | / |
|  |  | Allele 2 | ATGGGGTCGGCCTTTG**.....**GTAGTCCGGAGAGTGGTCCAGGAGCTGGACCAT | 38 |
|  |  | Allele 3 | ATGGGGTCGGCCTTT**T**AGCGGGTAGTCCGGAGAGTGGTCCAGGAGCTGGACCAT | 4 |
| *GSDMD^-/-^ XPR1^KSC/KSC^* cl2 | *XPR1* | WT | TCATTTCTTTCTGTAGAATCTGAATTTTACAGGGTTTCGAAAAATCCTGAAAAAGCATGACAAGATCCTGGAAACATCTCGTGGAGCAGATTGGCGAGTGGCTCACGTAGAG | 696 |
|  |  | HDR donor template | TCATTTCTTTCTGTAGAATCTGAATTTTACAGGGTTTCGA**GC**AATCCTG**GC**AAAGCATGAC**GC**GATCCTGGAAACATCTCGTGGAGCAGATTGGCGAGTGGCTCACGTAGAG | 696 |
|  | *GSDMD* | WT | ATGGGGTCGGCCTTTGAGCGGGTAGTCCGGAGAGTGGTCCAGGAGCTGGACCAT | 484 |
|  |  | Allele 1 | ATGGGGTCGGCCTTTG**..**CGGGTAGTCCGGAGAGTGGTCCAGGAGCTGGACCAT | 39 |
|  |  | Allele 2 | ATGGGGTCGGCCTTTGA**.**CGGGTAGTCCGGAGAGTGGTCCAGGAGCTGGACCAT | 6 |
